# PKC-dependent MYRF dysregulation links Aβ pathology to oligodendrocyte, myelin and cognitive alterations in Alzheimer’s disease

**DOI:** 10.64898/2025.12.09.693299

**Authors:** Uxue Balantzategi, Adhara Gaminde-Blasco, Neus Alcañiz, Juanma Ramirez, Rodrigo Senovilla-Ganzo, Nagore Hernández-Pinedo, Mario Fernandez-Ballester, Federico N. Soria, Alerie G. de la Fuente, Fernando García-Moreno, Estibaliz Capetillo-Zarate, Ugo Mayor, Edgar Soria-Gómez, José L. Zugaza, Elena Alberdi

## Abstract

Alzheimer’s disease (AD) is characterized not only by neuronal loss and synaptic dysfunction but also by emerging evidence of oligodendrocyte and myelin pathology. Amyloid-β (Aβ), a hallmark of AD, disrupts oligodendrocyte homeostasis through mechanisms that remain poorly understood. Here, we investigated the role of the myelin regulatory factor (MYRF), a transcription factor essential for oligodendrocyte maturation, in AD-related glial dysfunction. Using the 3xTg-AD mouse model, we observed aberrantly induced maturation dynamics and reduced oligodendrocyte lineage cell density at 12 months in the dentate gyrus, accompanied by increased MYRF expression. Interestingly, sustained MYRF overexpression was found to be toxic for oligodendrocytes *in vitro*. Moreover, *in vitro* and *in vivo* experiments further demonstrated that Aβ exposure elevates MYRF protein levels and enhances its transcriptional activity, pointing to post-translational regulation. Mechanistically, Aβ impaired GSK3-dependent phosphorylation and Fbxw7-mediated ubiquitination of MYRF, prolonging N-MYRF stability, an effect prevented by PKC inhibition *in vitro*. Intracerebroventricular infusion of PKC inhibitor Gö6983 normalized MYRF levels, restored oligodendrocyte populations and myelin integrity, and improved hippocampal-dependent spatial learning in 3xTg-AD mice, with locomotor activity and anxiety-like behavior remaining unaffected. Together, these findings identify MYRF dysregulation as a mechanistic link between Aβ/PKC signaling and oligodendrocyte pathology, and highlight PKC inhibition as a potential strategy to restore oligodendroglial function and cognition in AD.

## INTRODUCTION

Alzheimer’s disease (AD) is the most prevalent form of dementia, affecting over 50 million people worldwide. Traditionally, AD has been defined as a gray matter disorder characterized by progressive neuronal loss, synaptic dysfunction, and hallmark histopathologies including extracellular amyloid-β (Aβ) plaques and intracellular neurofibrillary tangles composed of hyperphosphorylated tau (Selkoe & Hardy, 2016). However, recent evidence highlights that white matter integrity and oligodendrocyte (OL) lineage cells also play critical roles in disease progression (Festa et al., 2024; Kedia & Simons, 2025). This paradigm shift has opened new avenues to understand how glial dysfunction contributes to cognitive decline in AD.

The triple transgenic 3xTg-AD mouse model, which harbors the human APP_Swe_, Tau_P301L_, and PS1_M146V_ mutations, is considered one of the most comprehensive animal models of the disease. It recapitulates both amyloid and tau pathology, as well as age-dependent cognitive impairments resembling those observed in patients (Oddo et al., 2003). Importantly, 3xTg-AD mice also exhibit alterations in myelin and OL biology (Qiu et al., 2023; Vanzulli et al., 2020), making them a valuable tool to study glial contributions to AD pathogenesis.

Accumulating evidence from human post-mortem samples and AD mouse models indicates disruption of myelin microstructure (Bartzokis et al., 2003; Han et al., 2002; Wu et al., 2017) and abnormalities in OL lineage dynamics, including impaired oligodendrogenesis (Behrendt et al., 2013; Dong et al., 2018) and defective maturation (Desai et al., 2010). Notably, Aβ itself has been shown to influence OL biology: previous work from our group demonstrated that exposure to Aβ precociously drives OL differentiation *in vitro* (Quintela-López et al., 2019) and in an *in vivo* zebrafish model (Balantzategi et al., 2025). These findings suggest that OL dysfunction may actively contribute to AD pathology by impairing myelination, axonal conduction, and neuronal network stability. Nevertheless, discrepancies across studies highlight the need for further investigation into the underlying molecular events.

During development and in adulthood, oligodendrocyte precursor cells (OPCs) undergo a tightly regulated differentiation program, progressing through immature pre-myelinating stages before becoming mature myelinating OLs, each stage defined by distinct molecular markers and morphological changes (Baumann & Pham-Dinh, 2001). OLs are essential for the formation and maintenance of myelin in the CNS (Simons & Nave, 2016). In the hippocampus, a region central to learning and memory, ongoing adult oligodendrogenesis supports neuronal plasticity and cognitive processes, and its disruption has been linked to neurodegenerative pathology (Hu et al., 2025; Steadman et al., 2020).

A key transcriptional regulator of OL maturation is the myelin regulatory factor (MYRF), a membrane-bound transcription factor essential for terminal OL differentiation and CNS myelination (Aprato et al., 2020; Bujalka et al., 2013; Emery et al., 2009). MYRF is synthesized in the endoplasmic reticulum (ER), undergoes homo-trimerization, and this full-length form (fl-MYRF) is then autoproteolytically cleaved. While the C-terminal fragment stays in the ER, the N-terminal fragment (N-MYRF) translocates to the nucleus, where it drives transcription of myelin genes, such as myelin basic protein (MBP), myelin-associated glycoprotein (MAG), and Myelin Oligodendrocyte Glycoprotein (MOG). MYRF degradation is regulated by GSK3-driven phosphorylation and ubiquitin-dependent degradation pathways (Nakayama et al., 2018). Yet, most of this knowledge derives from non-glial systems, and its regulation in OLs remains poorly understood. Given the central role of MYRF in OLs and myelin biology, it is reasonable to think that its dysregulation could have profound consequences in pathology; however, its implication in AD remains unknown.

Here, we report that in the dentate gyrus of 12-month-old 3xTg-AD mice, OL maturation is aberrantly enhanced and OL density is reduced, accompanied by increased MYRF expression. Moreover, *in vitro* and *in vivo* experiments show that Aβ exposure upregulates MYRF protein levels and enhances its transcriptional activity. Interestingly, sustained MYRF overexpression compromised OL viability *in vitro*. Mechanistic analyses revealed that Aβ impairs GSK3-dependent signaling and Fbxw7-derived ubiquitination of MYRF, thereby impairing its degradation and prolonging stability, in a PKC-dependent manner. Finally, continuous intracerebroventricular infusion of the PKC inhibitor Gö6983 normalized MYRF levels, restored OL populations and myelin integrity, and improved hippocampal-dependent spatial learning in 3xTg-AD mice, without affecting locomotion or anxiety-like behavior. Together, we propose MYRF dysregulation as an important contributor to AD pathology, and identify PKC signaling as a mechanistic link between Aβ and OL, myelin, and cognitive alterations in the disease.

## MATERIAL AND METHODS

### Animals

Experiments were performed in Sprague Dawley rat pups, C57BL6/J mice, and the triple transgenic mouse model of Alzheimeŕs disease (3xTg-AD), which harbors the APP_Swe_, Tau_P301L_, and PS1_M146V_ mutations (Oddo et al., 2003); wild-type B6129SF2/J mice were used as controls. Animals were housed in standard conditions with a 12-h light/dark cycle and *ad libitum* food and water. Procedures were conducted under the supervision and approved by the Ethics Committees of the University of the Basque Country (EHU) in accordance with the European Communities Council Directive 2010/63/EU. All possible efforts were made to minimize animal suffering and reduce the number of animals used.

### EdU labeling of proliferating cells

EdU labeling was performed with the Click-iT^TM^ EdU Alexa Fluor 647 Imaging Kit (Invitrogen), following manufacturer’s instructions. WT and 3xTg-AD mice (11 months, n = 4 per group) received 5’-ethynyl-2’-deoxyuridine (EdU; 0.2 mg/ml; Invitrogen) in drinking water for 30 days (Young et al., 2013); water was renewed every 48 h. After treatment, animals were perfused and brains sectioned. EdU was visualized incubating the brain slices with the Click-iT^TM^ reaction cocktail prior to immunofluorescence.

### Preparation of amyloid-β oligomers

Oligomeric amyloid-β (Aβ_1-42_) was prepared as previously described (Dahlgren et al., 2002). Briefly, synthetic Aβ_1-42_ peptide (Bachem, Germany) was initially dissolved in hexafluoroisopropanol (HFIP; Sigma-Aldrich) to 1 mM, aliquoted, and dried under vacuum. The resulting peptide films were stored desiccated at -80°C. For aggregation, peptide was resuspended in dry dimethylsulfoxide (DMSO; Sigma-Aldrich) at 5 mM and diluted in Ham’s F-12 medium (PromoCell) to a final concentration of 100 μM. Peptide solution was incubated at 4 °C for 24 h to obtain oligomeric species. Control cells were treated with vehicle (DMSO + Ham’s F-12).

### Intrahippocampal Aβ injections

Adult male C57BL6/J mice (3-4 months old) were randomly assigned to treatment groups (n = 5 per group) and anesthetized with ketamine hydrochloride (80 mg/kg) and xylazine (10 mg/kg). Animals were secured in a stereotaxic apparatus, and 3 µL of either vehicle (17% DMSO + 83% Ham’s F12) or Aβ (10 µM) was injected into the right dentate gyrus (coordinates: AP -2.2 mm, ML +1.5 mm, DV -2.0 mm) using a Hamilton microsyringe. The needle was left in place for 5 min to minimize reflux.

7 days after injection, mice were re-anesthetized with ketamine/xylazine and perfused transcardially with 30 mL of phosphate buffer (PB) followed by 30 mL of 4% paraformaldehyde (PFA) in 0.4 M PBS (pH 7.5). Brains were extracted and post-fixed in the same fixative solution for 4 h at room temperature (RT), transferred to 30% sucrose in 0.1 M PBS (pH 7.5) at 4 °C, and stored in cryoprotectant solution (30% ethylene glycol, 30% glycerol and 0.1 M PBS in dH_2_O) at -20 °C.

### Cell cultures

#### Primary oligodendrocyte culture

Highly enriched OPCs were obtained from mixed glial cultures prepared from the forebrain cortices of newborn (P_0_-P_2_) Sprague-Dawley rats, following established protocols (McCarthy & De Vellis, 1980) with modifications (Sánchez-Gómez et al., 2018). Briefly, cortices were dissected, meninges removed, and tissue enzymatically digested in Ca^2+^/Mg^2+^-free Hank’s balanced salt solution (HBSS-/-) containing 0.25% trypsin and 0.004% deoxyribonuclease (DNase; Sigma-Aldrich) at 37 °C for 15 min. The reaction was stopped with Iscove’s modified Dulbecco’s medium (IMDM; Gibco) supplemented with 10% fetal bovine serum (FBS HyClone). Cells were centrifuged (1,000 x g, 5 min), mechanically dissociated, and plated in poly-D-lysine (PDL; 1 μg/ml; Sigma-Aldrich)-coated T75 flasks. Mixed glial cultures were maintained at 37 °C, 5% CO_2_ until they were confluent (8-10 days).

Microglia were removed by shaking the flasks on a rotatory shaker (400 rpm, 1 h, 37 °C), and OPCs were isolated by further shaking (200 rpm, 18 h, 37 °C). OPCs were filtered through a 10 μm nylon mesh, pre-plated for 45 min in 100 mm coated Petri dishes (ThermoFisher Scientific) to eliminate residual microglia, and resuspended in chemically defined SATO medium (DMEM supplemented with 4.5 g/l glucose, 0.11 g/l sodium pyruvate, and growth/differentiation factors). All experiments were performed under SATO medium supplemented for differentiation (SATO^+^), except for MYRF overexpression assays, which were conducted in proliferation medium (SATO^-^) to ensure differentiation was driven by MYRF, rather than by the differentiating factors in the medium. Cells were seeded onto PDL-coated plates, with or without coverslips depending on experimental requirements, at densities of 1 x10^4^ – 1 x10^6^ cells/well, and maintained at 37 °C, 5% CO_2_ for 3 days. At 3-4 DIV, 88% of the cells were MBP^+^ OLs (Sánchez-Gómez et al., 2018).

#### HEK293T cells

Human embryonic kidney 293T (HEK293T) cells were maintained in Dulbecco’s modified Eagle medium (DMEM; Gibco) supplemented with 10% FBS (Gibco) and 1% penicillin/streptomycin (P/S; Gibco) at 37 °C, 5% CO_2_. Cells were passaged every 3-4 days to preserve 80-90% confluence. For reseeding, cells were detached with 0.25% trypsin-EDTA (Sigma-Aldrich) at 37 °C for 2 min and plated at densities of 1 x10^4^ - 1×10^6^ per well, according to experimental requirements.

### DNA plasmids and siRNAs

#### Gene expression via DNA plasmids

Exogenous MYRF expression was achieved using pcDNA3-Myc-Myrf-Flag (Bujalka et al., 2013) and GFP-C11orf9 (N-terminal GFP-tagged MYRF; C11orf9 is the former MYRF nomenclature) (Li et al., 2013). For luciferase assays, reporter constructs under the control of *Mbp* (pGL3-MBP), *Mag* (pGL3-MAG) (Bujalka et al., 2013), and *Myrf* (pGL4.10_myrfprom_ECR9) (Hornig et al., 2013) enhancers were used. Ubiquitin studies employed FLAG-tagged ubiquitin (pCDNA3.1-FLAG-Ub) (Ramirez et al., 2018), HA-tagged ubiquitin plasmids (WT; pRK5-HA-Ub-K48 (#17605, Addgene); and pRK5-HA-Ub-K63 (#17606, Addgene)), untagged human Parkin (Martinez et al., 2017), and Myc-tagged human Fbxw7 (pCMV3-Myc-FBXW7-t2; #HG29625-NM, Sino Biological).

Plasmid DNA was purified using QIAGEN Plasmid Maxi (Qiagen) or NZYMaxiprep (NZYtech) kits, and concentrations were determined with a Nanodrop 2000 spectrophotometer (ThermoFisher Scientific). Primary OLs were transfected by electroporation with the Amaxa^TM^ Basic Nucleofector^TM^ Kit for Primary Mammalian Glial cells (Lonza), whereas HEK293T cells were transfected by lipofection using Lipofectamine 3000^TM^ (Invitrogen), following manufacturers’ instructions. A GFP construct was routinely co-transfected as a positive control for transfection efficiency, which was monitored by fluorescence microscopy (EVOS^TM^ FL Digital Inverted Microscope; Invitrogen).

#### siRNA transfections

For *Myrf* knockdown, primary OLs were nucleofected with 100 pmol of either scrambled control siRNA (ON-TARGETplus Non-targeting Pool, #D-001810-10-05) or rat *Myrf* targeting siRNA (ON-TARGETplus Rat Myrf siRNA, #L-085053-02-05) (Dharmacon, HorizonDiscovery) using the Amaxa^TM^ Basic Nucleofector^TM^ Kit for Primary Mammalian Glial cells (Lonza), following the manufacturer’s instructions. Transfected cells were seeded onto PDL-coated 24-well plates with coverslips at densities 1 x 10^5^ cells/well for western blot or 7.5×10^4^ cells/well for luciferase assays.

### Immunofluorescence

#### Immunocytochemistry of primary oligodendrocytes

Primary cultured OLs were exposed to vehicle or Aβ as indicated. Cells were fixed in 4% PFA with 4% sucrose for 15 min at RT, rinsed in PBS, and permeabilized/blocked for 1 h in PBS containing 4% normal goat serum (NGS; Palex) and 0.1% Triton X-100 (Sigma-Aldrich). Cells were incubated overnight (O/N) at 4 °C with primary antibodies against Myc tag (ms IgG_1_, 1:500; #05-724, Millipore) and Olig2 (ms IgG_2A_, 1:1,000; #MABN50, Millipore) diluted in blocking solution. After three washes in PBS, samples were incubated with Alexa Fluor-conjugated secondary antibodies (1:500; Invitrogen) in blocking solution for 1 h at RT. Nuclei were counterstained with DAPI (4 μg/ml; Sigma-Aldrich) for 10 min, and coverslips were mounted with Fluoromount-G^®^ (SouthernBiotech). Images were acquired on a Leica TCS STED CW SP8X confocal microscope (Leica) using a 63X oil-immersion objective. Identical acquisition settings were applied across conditions, and image analysis was performed with ImageJ/Fiji (NIH).

#### Immunohistochemistry of mouse brain tissue

Mice were anesthetized by intraperitoneal injection of pentobarbital and perfused with 4% PFA (Electron Microscopy Sciences) in PBS. Brains were extracted, post-fixed for 4 h at RT, rinsed in PBS, and sectioned into 40 μm coronal slices using a vibratome (Leica microsystems). Free-floating sections were washed in 0.1 M PB and, when required for nuclear epitope accessibility, subjected to antigen retrieval in R-Universal Epitope Recovery Buffer (Aptum) for 5 min at 95 °C, followed by 5 min incubation at RT. Sections were rinsed three times in PBS and permeabilized/blocked in PBS containing 10% NGS and 0.1% Triton X-100 for 30 – 60 min at RT. Samples were incubated O/N at 4 °C with primary antibodies in blocking solution: anti-Olig2 (ms IgG_2A_, 1:500; #MABN50, Millipore), anti-APC (CC1) (ms IgG_2B_, 1:200; #OP80, Millipore), anti-PDGFRα (gt, 1:500; #AF1062, Biotechne), anti-MYRF (rb, 1:500; #A16355, ABclonal), anti-Caspr (ms, 1:500; #K65/35, Neuromab), and anti-Na_v_1.6 (rb, 1:1,000; #ASC-009 Alomone). After three washes in PBS, sections were incubated with Alexa Fluor-conjugated secondary antibodies (1:500) for 1 h at RT, counterstained with DAPI for 10 min, and washed again before mounting with Fluoromount-G^®^ mounting medium. Z-stack images of the dentate gyrus were acquired on a Leica TCS STED CW SP8X confocal microscope, and image analysis was performed with ImageJ/Fiji software (NIH).

### Protein preparation and western blot analysis

#### Protein preparation from primary oligodendrocytes

After treatments, OLs were washed twice in PBS and scraped in sodium dodecyl sulfate (SDS) sample buffer (62.5 mM Tris pH 6.8, 10% glycerol, 2% SDS, 0.002% bromophenol blue, 5.7% β-mercaptoethanol in dH_2_O) on ice. At least two wells per condition (8 x10^4^ – 1 x10^5^ cells/well) were pooled.

#### Protein preparation from mouse hippocampi

Mice were anesthetized by intraperitoneal injection of pentobarbital and perfused with saline (0.9% NaCl in H_2_O). Hippocampi were dissected, snap-frozen on dry ice, and stored at -80 °C. Tissue samples were resuspended in 50 μl RIPA lysis buffer (ThermoFisher) supplemented with protease and phosphatase inhibitors, homogenized with a douncer, and adjusted to 200 μl with additional RIPA buffer. Tissue homogenates were sonicated (25 pulses, 80% amplitude, 0.6 cycle; Labsonic M, Sartorius), centrifuged (2,000 x g, 10 min, 4°C), and supernatants collected. Protein concentration was determined by the Bradford assay (Bio-Rad), and samples were prepared with SDS sample buffer.

#### Western blot

Samples were boiled at 95 °C for 5 min, size-separated by electrophoresis in 4-20% (Any KD) polyacrylamide-SDS Criterion TGX Precast gels (Bio-Rad), and transferred to Trans-Blot^®^ Turbo™ Midi Nitrocellulose Transfer Packs (Bio-Rad). Membranes were blocked in 5% BSA in TBST for 1 h at RT, and incubated O/N at 4 °C with the following primary antibodies diluted in blocking solution: anti-Myc tag (ms, 1:2,000; #05-724, Millipore), anti-FLAG tag (rb, 1:1,000; #2368, Cell Signaling), anti-MYRF (rb, 1:1,000; #ABN45, Millipore), anti-MBP (ms, 1:1,000; #SMI-99, BioLegend), anti_-_ p-GSK3α/β (Ser21/9) (rb, 1:1,000; #9331, Cell Signaling), anti-GSK3α/β (ms, 1:500; #sc-7291, Santa Cruz), and anti-GAPDH (ms, 1:5,000; #MAB374, Millipore). After three washes in TBST, membranes were incubated for 1 h at RT with horseradish peroxidase (HRP)-conjugated (1:5,000) or fluorescent secondary antibodies (1:5,000), washed three times in TBST and once in TBS. HRP signals were developed using enhanced chemiluminescence (NZY standard or advanced ECL; NZYtech). Protein bands were imaged with a ChemiDoc XRS Imaging System (Bio-Rad), and quantified using Image Lab^®^ 6.0.1 software (Bio-Rad). When required, membranes were stripped with Restore Western Blot Stripping Buffer (ThermoFisher Scientific) following the manufacturer’s instructions.

### Dual-luciferase assays

Primary cultured OLs were co-transfected with *firefly* luciferase reporter construct (pGL3-Basic, pGL3-Control, pGL3-MBP, or pGL3-MAG) together with a constitutive *Renilla* luciferase construct (ratio 1:10) as internal control. pGL3-Basic (promoterless) was used as a negative control, whereas pGL3-Control (SV40 promoter, lacking *Myrf*-responsive elements) served as reference reporter, with its activity arbitrarily set to 1. Cells were additionally co-transfected with control or *Myrf* siRNAs, as previously described. Transfected OLs were seeded in PDL-coated 24-well plates in triplicate at a density of 7.5×10^4^ cells/well.

At 72 h post-transfection (3 DIV), cells were lysed in Passive Lysis Buffer (1X PLB, Promega), and luciferase activity was measured using the Dual-Luciferase® Reporter Assay System (Promega) according to the manufacturer’s instructions. For each condition, 10 μl of cell lysate was transferred to white opaque 96-well plates. *Firefly* luciferase activity was measured following addition of 50 μl Luciferase Assay Reagent II (LAR II, Promega) using a Synergy H4 Hybrid Reader (BioTek Instruments). Subsequently, 50 μl Stop & Glo Reagent (Promega) was added to each well, and *Renilla* luciferase activity was recorded. All luminescence values were normalized to *Renilla* activity and further corrected for protein concentration determined by the Lowry assay (Bio-Rad).

### Cell viability assays

Primary OLs were electroporated with an empty plasmid or the double-tagged MYRF construct, and seeded in 48-well plates (2 x10^4^ cells/well, triplicates) in SATO-medium. Cells were incubated with Calcein-AM (1 μM; Invitrogen) and NucBlue^TM^ Live ReadyProbes Reagent (2 drops/ml; Invitrogen) at 3, 6, and 10 DIV for 30 min at 37°C. After two PBS washes, cells were imaged in an EVOS fluorescence microscope (Invitrogen), and fluorescence quantified in a CLARIOstar^®^ Plus microplate reader (BMG Labtech). Excitation/emission settings: 485/528 nm (Calcein-AM) and 360/460 nm (NucBlue). Viability was expressed as Calcein-AM (living cells)/NucBlue (total cells) ratio.

### Cycloheximide chase assay

Protein stability of N-MYRF was assessed by cycloheximide (CHX) chase in primary cultured OLs. Cells (1 x10^5^/well, duplicates) were seeded in PDL-coated 24-well plates and maintained in SATO+ medium for 3 DIV. CHX (100 µg/ml; Sigma-Aldrich) was added directly to the culture medium, and cells were harvested at the indicated time points. Aβ (1 µM) was added 1 h prior to CHX exposure. As a positive control for inhibition of MYRF degradation, cells were cultured with the GSK-3 inhibitor AR-A014418 (1 µM, 24 h). Cell lysates were prepared as described and analyzed by Western blotting using anti-MYRF (rb, 1:1,000; #ABN45, Millipore) and anti-GAPDH (ms, 1:5,000; #MAB374, Millipore). Degradation kinetics were expressed as the percentage of N-MYRF remaining at each chase time, with 100% representing baseline expression for each condition. Protein half-life was calculated using GraphPad Prism 8.2.1 software.

### Phosphoserine immunoprecipitation

HEK293T cells were seeded into 6-well plates at a density of 7.5 x10^5^ cells/well. The following day, at 70-90% confluence, cells were transfected with 2.5 µg GFP-MYRF plasmid per well using Lipofectamine 3000 (Invitrogen). Cells were incubated for 48 h at 37 °C with 5% CO_2_, and transfection efficiency was verified by GFP visualization using an EVOS fluorescence microscope (Invitrogen). Cells were then treated with vehicle, Aβ (1 µM; 15, 30, or 60 min), or the GSK3 inhibitor AR-A014418 (1 µM) as a positive control for inhibition of GSK3-mediated phosphorylation.

For immunoprecipitation, cells were washed twice with cold PBS, lysed in 100 μl supplemented RIPA lysis buffer (ThermoFisher Scientific), incubated on ice for 10 min, and centrifuged (12,000 × g, 5 min, 4 °C). One-tenth of the supernatant was saved as input, and the remaining was incubated with 1 μg agarose-conjugated mouse anti-phospho-serine (pSer) antibody (#A8076, Sigma-Aldrich) or mouse IgG control (#sc-2342 AC, Santa Cruz Biotechnology) pre-washed twice in RIPA. The lysate-antibody-beads complexes were rotated for 2 h at 4°C, centrifuged (2,000 x g, 2 min) to separate beads from the unbound material, and washed twice with RIPA and once with PBS. Finally, bound proteins were eluted in 50 μl 2x SDS sample buffer, by boiling the samples at 95°C for 5 min, followed by centrifugation (14,000 x g, 1 min). Both inputs and eluates were analyzed by Western blotting using anti-MYRF (rb, 1:1,000; #ABN45, Millipore) and anti-GAPDH (ms, 1:5,000; #MAB374, Millipore) antibodies.

### Ubiquitin analysis by GFP pull-down

HEK293T cells were seeded in 6-well plates at a density of 1 x 10^6^ cells per well. For ubiquitination level experiment, each well was co-transfected with 1 µg of GFP-MYRF, 1 µg FLAG-Ub, and 1 µg of either empty plasmid (ø), Myc-Fbxw7, or untagged Parkin. To assess the type of ubiquitination, cells were co-transfected with 1 µg of GFP-MYRF and 1 µg of Myc-Fbxw7, together with 1 µg WT, K48, or K63 HA-tagged ubiquitin plasmids. At 48 h post-transfection, cells were treated with Aβ (1 µM) or vehicle for 3 h before GFP pull-down.

Pull-down was performed as previously described (Elu et al., 2019). Briefly, cells were washed twice with PBS and lysed in 300 µl of lysis buffer (50 mM Tris–HCl pH 7.5, 150 mM NaCl, 1 mM EDTA, 0.5% Triton, 1× protease inhibitor cocktail (Roche Applied Science), and 50 mM N-ethylmaleimide (NEM; Sigma-Aldrich)). Lysates were centrifuged (16,000 x g, 10 min, 4 °C), and one-tenth of each supernatant was saved as input. The remaining supernatant was incubated with 20 µl/condition of GFPTrap-A agarose beads (GFP-beads; Chromotek GmbH) previously washed twice in dilution buffer (10 mM Tris-HCl pH 7.5, 150 mM NaCl, 0.5 mM EDTA, 1× protease inhibitor cocktail, 50 mM NEM). After incubation of 2.5 h at RT with gentle rotation, complexes were pelleted by centrifugation (2,700 x g, 2 min), washed once with dilution buffer, three times with washing buffer (8 M urea, 1% SDS in PBS), and once with 1% SDS in PBS. Bound proteins were eluted by boiling beads at 95 °C for 10 min in 25 µl elution buffer (250 mM Tris–HCl pH 7.5, 40% glycerol, 4% SDS, 0.2% bromophenol blue, 100 mM DTT), followed by centrifugation (16,000 x g, 2 min).

MYRF-linked ubiquitin was detected by Western blotting using anti-FLAG (rb, 1:1,000; #2368, Cell Signaling) or anti-HA (ms, 1:1,000; #3663, Sigma-Aldrich) antibodies, depending on the experiment. Additional primary antibodies included anti-GFP (ms, 1:1,000; #11814460001, Millipore), anti-Fbxw7 (ms, 1:1,000; #TA802869S, OriGene), anti-Parkin (ms, 1:1,000; #sc-32282, Santa Cruz), and anti-α-Tubulin (ms, 1:5,000; #ab7291, abcam).

### Continuous intracerebroventricular Gö6983 infusion

12-to 14-month-old WT and 3xTg-AD mice were used (n = 3-5 per treatment group). Osmotic minipumps (Model 1004, Alzet) were subcutaneously implanted and connected to an infusion headstage attached to the skull (Brain infusion kit II, Alzet), continuously delivering either vehicle (0.01% DMSO in saline) or Gö6983 (100 nM) at 0.11 µl/h for 28 days (Liu et al., 2019), following manufacturer’s instructions.

On the day of surgery, mice were anesthetized with isoflurane via a precision vaporizer, and both the scalp and implantation site were shaved and disinfected. Animals were secured in a stereotaxic frame, a midline sagittal incision was made to expose the skull, and a subcutaneous pocket was prepared in the midscapular region for pump placement. Using stereotaxic coordinates (AP -0.5 mm, ML +1 mm, DV -2 mm), a cranial hole was drilled to insert the cannula connected to the osmotic pump, enabling continuous infusion into the right lateral ventricle. The pump was positioned in the subcutaneous pocket, the headstage secured with dental cement, and the incision closed with sutures.

Postoperatively, mice received buprenorphine (0.3 mg/kg of animal) for analgesia. Animal well-being was monitored throughout the experiment, and body weight was recorded before, during, and after treatment. After 28 days, mice were anesthetized with pentobarbital (i.p.) and perfused with saline. Brains were dissected; right hemispheres (ipsilateral) were post-fixed in 4% PFA in PBS O/N at 4 °C for IHC, and left hippocampi (contralateral) were snap-frozen for protein extraction and Western blot analysis.

### Behavioral test: Barnes Maze

Mice carrying osmotic minipumps were allowed to recover for 2 weeks post-surgery and were subjected to behavioral testing during the last 14 days of treatment. Spatial learning and memory were assessed using the Barnes Maze (BM), a dry-land behavioral test widely employed to evaluate hippocampus-dependent cognition (Bach et al., 1995; Bird & Burgess, 2008; Pitts, 2018). The apparatus consisted of a circular white platform (150 cm diameter, 90 cm height) with 20 evenly spaced peripheral holes, one connected to an escape chamber. Each mouse was assigned a fixed escape hole location throughout the experiment, with distinct locations across animals. Bright illumination and open space served as aversive stimuli, motivating mice to find the escape box, while distal visual cues placed around the room provided spatial references (Rueda-Orozco et al., 2008).

Test was performed over 5 consecutive days, 4 trials/day. Mice were acclimated to the room for 20-30 min before each session. At trial start, animals were placed in the center of the maze inside an opaque cylinder for 5 s, then released. Each trial lasted max 4 min, followed by 1 min rest in the home cage while the maze was cleaned. On day 1, mice were guided to the target hole prior to the first trial, as a training. If a mouse failed to find the box within 4 min (non-successful), it was guided to it. Trials were recorded and analyzed using ANY-maze software (ANY-maze; Dublin, Ireland). Performance was quantified by latency to enter the target escape hole and by search strategy. A spatial strategy was defined when mice moved directly to the target or adjacent holes (hippocampal-dependent strategy).

### Statistical analysis

All statistical analyses were performed using GraphPad Prism 8.2.1 (GraphPad Software). Data are presented as mean ± S.E.M, with sample size (n) indicated in the figures by individual data points. Statistical comparisons between two groups were assessed using unpaired or paired two-tailed Student’s t-tests. For multiple group comparisons, one-way or two-way analysis of variance (ANOVA) was applied, followed by Dunnett’s, Tukey’s, or Sidak’s post hoc tests, as appropriate. Analyses were conducted using absolute values. Independent animals or separate experiments were considered biological replicates (n ≥ 3). Statistical significance was defined as: p<0.05 (* or #), p<0.01 (** or # #), p<0.001 (*** or # # #), and p<0.0001 (****or # # # #).

## RESULTS

### 3xTg-AD mice exhibit promoted oligodendrocyte maturation and lineage cell loss in the hippocampus

Given the evidence for OL dysfunction in AD, we began by assessing OL lineage dynamics in the hippocampus of 12-month-old 3xTg-AD mice. With that aim, at 11 months, WT and 3xTg-AD mice received EdU (0.2mg/ml) in drinking water for 30 days to label proliferating cells (**Fig. 1a**). Immunohistochemistry was performed on hippocampal sections (**Fig. 1c, d**), focusing on the hilus of the dentate gyrus, a region with prominent adult oligodendrogenesis and vulnerability to AD-related pathology (DeFlitch et al., 2022). Lineage cells were classified into OPCs (PDGFRα^+^), immature OLs (PDGFRα^-^CC1^-^Olig2^+^), and mature OLs (CC1^+^Olig2^+^) (**Fig. 1b**). 3xTg-AD mice showed fewer newly generated Olig2^+^EdU^+^ cells than WT (6 ± 0.73 x10^3^ cells/mm^3^ vs 11.43 ± 1.69 x10^3^ cells/mm^3^, respectively; **Fig. 1e**), but a significantly higher fraction of these newly generated cells matured into CC1^+^ OLs (42.79 ± 6.32% vs 19.84 ± 3.07%; **Fig. 1f**), indicating enhanced maturation efficiency. No sex differences were detected.

**Figure 1.**
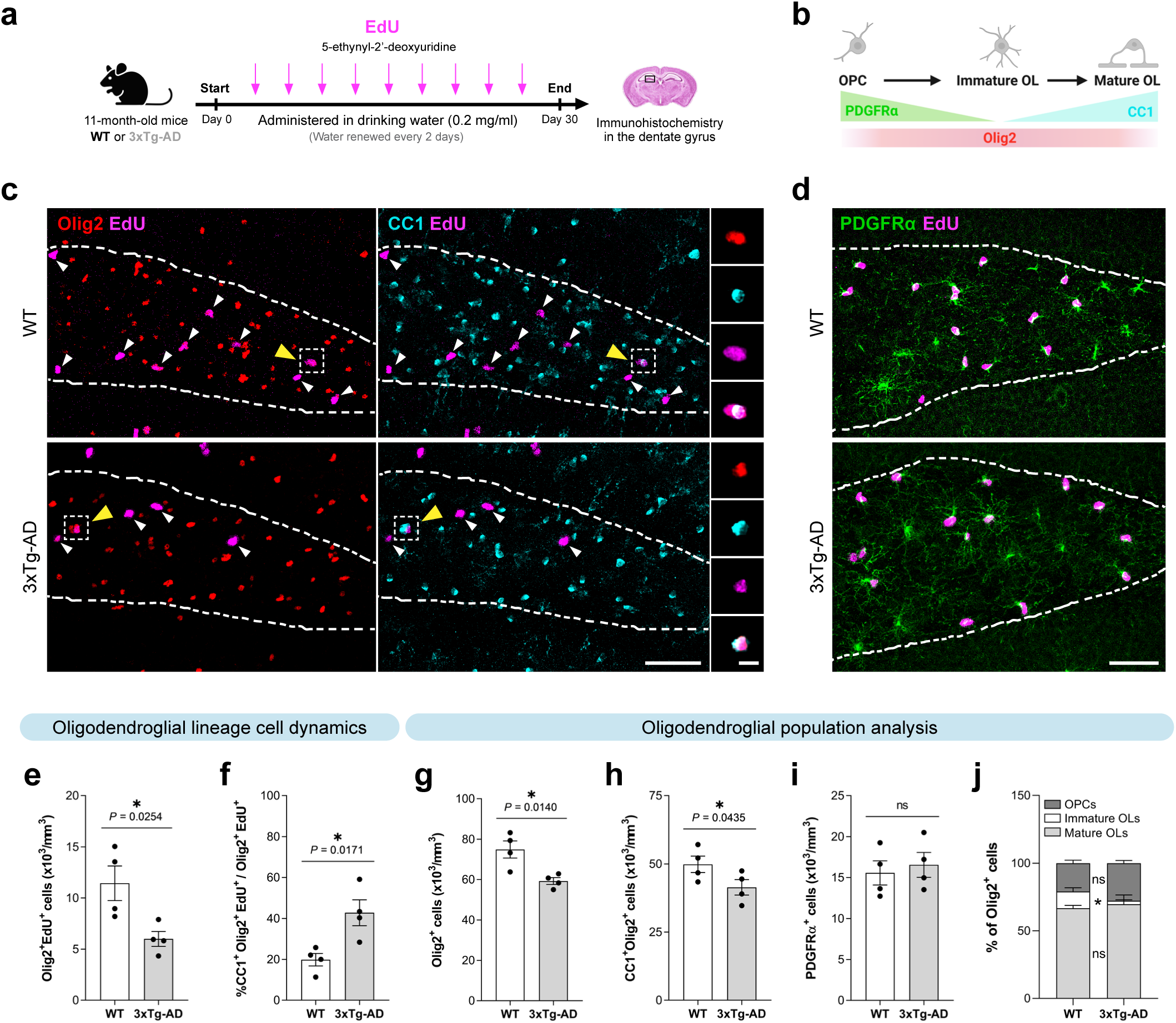
Oligodendroglial population is altered in the dentate gyrus of 12-month-old 3xTg-AD mice. (**a**) Experimental approach: 11-month-old WT and 3xTg-AD mice received EdU (0.2 mg/ml) in drinking water for 30 days, followed by brain dissection and IHC for analysis of oligodendroglial populations and dynamics. (**b**) Schematic representation of differentiating oligodendroglial subpopulations, and their specific markers. (**c**) Representative confocal images showing EdU (magenta), OL lineage marker Olig2 (red), mature OL marker CC1 (cyan), and (**d**) OPC marker PDGFRα (green) in the dentate gyrus; dotted white line delineates the hilus. White arrowheads indicate Olig2^+^EdU^+^ cells; yellow arrowheads mark newly generated mature OLs (CC1^+^Olig2^+^EdU^+^). Scale bar: 50 µm. (**e-f**) Quantification of oligodendroglial lineage cell dynamics: (**e**) Olig2^+^EdU^+^ cells (proliferating OLs) and (**f**) % CC1^+^Olig2^+^EdU^+^/Olig2^+^EdU^+^ (ratio of newly generated mature OLs). (**g-j**) Oligodendroglial population analysis: (**g**) total Olig2^+^ cells (all lineage cells), (**h**) CC1^+^Olig2^+^ cells (mature OLs), (**i**) PDGFRα^+^ cells (OPCs), and (**j**) relative fractions of oligodendroglial subpopulations (OPCs, immature OLs, and mature OLs). Data are presented as means ± S.E.M; dots represent individual animals. *p<0.05; statistical significance was determined by two-tailed unpaired t-test (**e**, **f**, **g**, **h**, **i**) or two-way ANOVA followed by Sidak’s *post-hoc* test (**j**), as appropriate.

Next, when examining total oligodendroglial populations, 3xTg-AD mice showed reduced total Olig2^+^ cell density (59.23 ± 1.68 x10^3^ cells/mm^3^) compared to WT (74.92 ± 4.25 x10^3^ cells/mm^3^) (**Fig. 1g**), as well as mature CC1^+^Olig2^+^ OLs (41.38 ± 2.83 x10^3^ cells/mm^3^ vs 49.84 ± 3.02 x10^3^ cells/mm^3^; **Fig. 1h**). In fact, some Olig2^+^ cells in the 3xTg-AD dentate gyrus displayed pyknotic nuclei and were occasionally engulfed by microglia (**Supplementary Fig. 1**), suggesting possible cell death by phagoptosis. In contrast, PDGFRα^+^ OPC density remained unchanged across genotypes (**Fig. 1i**). Oligodendroglial subpopulation analysis revealed a marked reduction in the relative fraction of immature OLs in 3xTg-AD mice (2.5 ± 4.31%) versus WT (12.46 ± 2.75%), while OPC and mature OL proportions were similar (**Fig. 1j**).

Together, these findings reveal subpopulation-specific effects of AD on OL lineage, with aberrantly promoted maturation and near depletion of immature cells, likely reflecting a maladaptive attempt to replace lost mature OLs in 3xTg-AD mice.

### MYRF is upregulated in oligodendrocytes from AD mouse models

Given the enhanced differentiation of newly generated OLs observed in 3xTg-AD mice, we next investigated whether this phenotype might be linked to alterations in a key transcriptional regulator of OL maturation: MYRF. First, we assessed MYRF expression in the dentate gyrus of 12-month-old WT and 3xTg-AD mice by immunofluorescence. Quantification revealed a significantly higher proportion of MYRF^+^ OLs in the 3xTg-AD mice (83.6 ± 0.92%) compared to age-matched WT controls (72.1 ± 4.1%) (**Fig. 2a, b**), consistent with the aberrant OL differentiation previously observed in this model.

**Figure 2.**
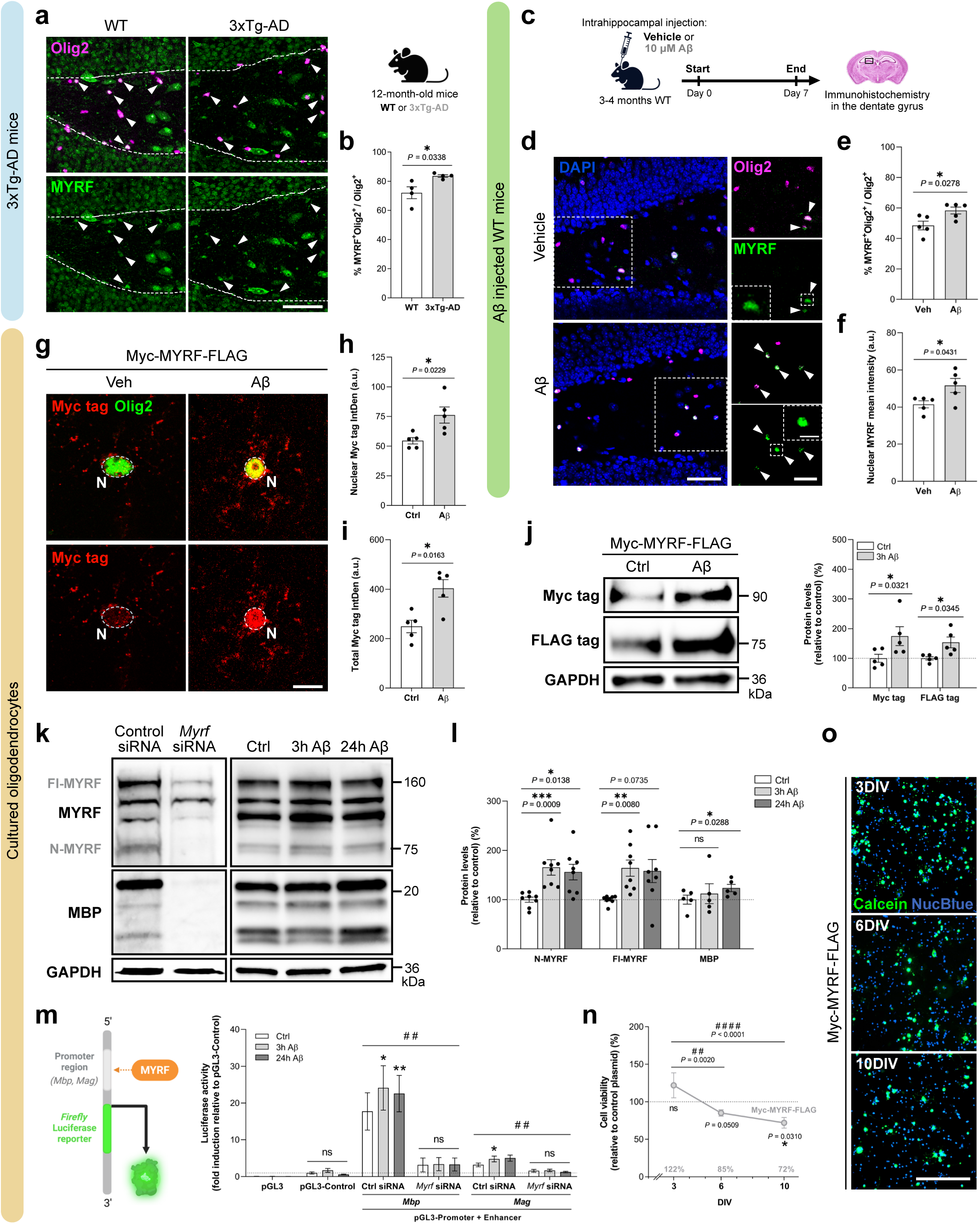
MYRF expression and transcriptional activity are increased in oligodendrocytes from Alzheimer’s disease models. (**a**) Immunofluorescence images of the dentate gyrus in 12-month-old WT and 3xTg-AD mice showing MYRF (green) and Olig2 (magenta). White arrows indicate MYRF^+^Olig2^+^ cells. Scale bar: 50 μm. (**b**) Quantification of MYRF^+^ OLs within the Olig2^+^ population. (**c-f**) Experimental design and MYRF analysis following intrahippocampal injection of vehicle or 10 µM Aβ into 3-4-month-old WT mice. (**d**) Immunohistochemistry of the dentate gyrus one week post-injection showing DAPI (blue), Olig2 (magenta), and MYRF (green). Arrows indicate MYRF^+^Olig2^+^ cells. Scale bars: 50 μm (overview), 25 μm and 10 μm (insets). (**e**) Percentage of MYRF^+^Olig2^+^ cells among total Olig2^+^ cells. (**f**) Mean nuclear MYRF intensity. (**g-j**) Primary OLs transfected with Myc-MYRF-FLAG and treated with vehicle or 1 µM Aβ for 3 h at 2 DIV. (**g**) Immunostaining for Olig2 (green) and Myc tag (red); dotted lines outline nuclei (N). Scale bar: 10 µm. (**h-i**) Quantification of (**h**) nuclear and (**i**) total Myc tag signal. (**j**) Western blot and quantification of cleaved N- and C-terminal MYRF (anti-Myc tag and -FLAG tag, respectively), normalized to GAPDH and expressed as percentage relative to vehicle. (**k**) Western blot of OPCs cultured in differentiation media for 3 DIV, either transfected with scramble (control siRNA) or *Myrf*-targeting siRNA, or treated with vehicle or Aβ for 3 or 24 h. Blots show full-length MYRF (Fl-MYRF; ∼160 kDa), cleaved N-terminal MYRF (N-MYRF; ∼75 kDa), MBP, and GAPDH. (**l**) Quantification of endogenous N-MYRF, Fl-MYRF, and MBP normalized to GAPDH and expressed as percentage relative to vehicle. (**m**) Luciferase reporter assay in 3 DIV OLs co-transfected with *Mbp* or *Mag* luciferase constructs, and either scrambled or *Myrf* siRNAs, and treated with vehicle or Aβ. Activity is shown as fold change relative to pGL3-Control plasmid. n = 3-6. (**n**) Calcein-AM viability assay in OLs overexpressing MYRF (Myc-MYRF-FLAG) for 3, 6, or 10 DIV, normalized to empty vector-transfected cells. (**o**) Representative fluorescence images of Calcein^+^ (live; green) and NucBlue^+^ (total; blue) cells. Scale bar: 200 µm. n^3^ ^DIV^ = 9; n^6^ ^DIV^ and n^10^ ^DIV^ = 6. Data are presented as mean ±S.E.M; dots represent independent experiments or animals. *p<0.05, **p<0.01, ***p<0.001 between treatments or plasmids (**n**); # #p<0.01 between siRNAs (**m**) or across time (**n**). Statistical significance was assessed by two-tailed unpaired (**b**, **e**, **f**) or paired (**h**, **i**, **j**) t-test, one-way ANOVA followed by Dunnett’s *post-hoc* test (**l**), or two-way ANOVA with Sidak’s (**m**, **n**) or Dunnett’s (**m**) *post-hoc* tests.

Since 3xTg-AD mice progressively accumulate Aβ in the hippocampus from early disease stages (Oddo et al., 2003), and Aβ has been linked to myelin disruption and OL dysregulation (Balantzategi et al., 2025; Ferreira et al., 2020; Quintela-López et al., 2019), we next asked whether Aβ contributes to MYRF upregulation. To test this, young adult WT mice (3-4 months old) received intrahippocampal injections of 10 µM Aβ or vehicle targeting the dentate gyrus, and brains were collected one week later for immunohistochemistry (**Fig. 2c**). Aβ-injected mice showed a significantly higher MYRF^+^Olig2^+^/Olig2^+^ ratio (58.3 ± 2.27%) than vehicle-treated controls (48.49 ± 2.87%) (**Fig. 2d, e**), indicating that Aβ promotes MYRF expression *in vivo*. Because MYRF becomes transcriptionally active upon nuclear translocation of its N-terminal fragment, we quantified nuclear MYRF intensity in those MYRF^+^Olig2^+^ cells. Aβ-treated mice showed a modest but significant increase in nuclear MYRF signal (51.62 ± 3.76 a.u.) compared to controls (41.46 ± 1.93 a.u) (**Fig. 2d, f**), suggesting enhanced MYRF activation following Aβ exposure.

Together, these findings indicate that MYRF is upregulated and potentially activated in OLs from both 3xTg-AD mice and WT mice exposed to Aβ, implicating MYRF in the altered dynamics of OL maturation in AD.

### Aβ increases MYRF expression and transcriptional activity in cultured oligodendrocytes

To further investigate how Aβ regulates MYRF, we moved to *in vitro* models. Using a dual-tagged MYRF construct (Myc-MYRF-FLAG), we characterized its cleavage products, their cellular localization, and molecular weights in HEK293T cells (**Supplementary Fig. 2a, b**), and confirmed in primary rat OLs that MYRF overexpression drives robust MBP induction, consistent with its well-established pro-myelinating role (**Supplementary Fig. 2c-e**).

To test whether Aβ modulates MYRF protein levels, primary OLs were transfected with MYRF construct and treated with 1 µM Aβ or vehicle for 3 hours. Immunostaining revealed a significant increase in both nuclear (vehicle: 54.53 ± 2.72 a.u. vs Aβ: 76.18 ± 6.77 a.u.; **Fig. 2g, h**) and total MYRF signal (vehicle: 249.11 ± 25.15 a.u. vs Aβ: 403.67 ± 35.29 a.u.; **Fig. 2g, i**). Western blot confirmed elevated levels of both cleaved N- (detected by Myc tag: 174.64 ± 31.87%) and C-terminal (FLAG tag: 153.57 ± 18.16%, relative to 100% of control; **Fig. 2j**) fragments upon Aβ treatment, indicating that Aβ promotes a global increase in MYRF protein without selectively targeting specific processing products.

To determine whether Aβ similarly affects endogenous MYRF, we optimized detection using RNA interference. Primary OLs transfected with *Myrf*-targeting siRNA (*Myrf* siRNA) showed markedly reduced MYRF protein levels and enabled clear identification of specific MYRF bands detected by the anti-MYRF antibody (∼75kDa for cleaved N-MYRF and ∼160kDa for full-length MYRF; **Fig. 2k**). Knockdown of *Myrf* also reduced MBP expression, underscoring MYRF’s essential role in OL maturation and myelin gene induction (Emery et al., 2009).

With this validated system, 3 DIV OLs were treated with vehicle or Aβ for 3 or 24 hours. Western blot revealed that Aβ significantly increased both N-MYRF (165.37 ± 15.69% at 3 h; 155.84 ± 15.89% at 24 h) and fl-MYRF (164.03 ± 16.48% at 3 h; 158.06 ± 23.5% at 24 h, relative to control), as well as MBP at 24 h (123.52 ± 7.42%; **Fig. 2k, l**), indicating that Aβ enhances MYRF levels and promotes downstream myelin protein expression.

Next, to assess whether Aβ also increases MYRF’s transcriptional activity, we performed dual-luciferase assays in primary OLs with *Mbp* and *Mag* enhancer luciferase constructs (Bujalka et al., 2013). As expected, these enhancers induced luciferase activity ∼18- and ∼3-fold, respectively, compared to pGL3-Control, which was significantly reduced to 6- and 1.6-fold, respectively, upon *Myrf* silencing. Interestingly, Aβ significantly increased *Mbp* (from 17.74 ± 5.07 to 24.12 ± 6.03 fold at 3 h; 22.58 ± 4.49 fold at 24 h) and *Mag* (from 3.2 ± 0.49 to 4.79 ± 0.78 fold at 3 h) enhancer-driven luciferase activity, an effect abolished by *Myrf* silencing (**Fig. 2m**), confirming that Aβ-induced transcriptional enhancement was MYRF-dependent. No changes were observed in pGL3-Control activity, confirming specificity.

Together, these data demonstrate that Aβ increases both the abundance and transcriptional activity of MYRF in OLs, supporting a model in which MYRF acts as a potential key mediator of Aβ-induced changes in oligodendroglial maturation in early AD.

### Prolonged MYRF overexpression impairs oligodendrocyte viability

While MYRF is selectively upregulated during OL differentiation and is essential for myelin gene expression (Emery et al., 2009), little is known about the potential adverse consequences of its sustained overexpression on OL survival. To address this, we assessed the effects of prolonged MYRF overexpression in primary cultured OLs using Calcein-AM viability assays at 3, 6, and 10 DIV. At 3 DIV, MYRF overexpression did not impair viability and even showed a non-significant trend toward increased survival (121.91 ± 16.73% vs. 100% in control cells). However, intriguingly, a progressive reduction in viability was observed over time in MYRF-overexpressing OLs compared to empty vector-transfected control cells (84.99 ± 4.29% at 6 DIV; 71.9 ± 7.05% at 10 DIV; **Fig. 2n**). Fluorescence images further illustrated this temporal decline in Calcein^+^ cells (**Fig. 2o**).

Overall, this data suggest that while transient MYRF expression promotes OL maturation, sustained and excessive MYRF levels compromise cell survival. These findings highlight that MYRF requires tight temporal regulation, and raises the possibility that its dysregulation in AD may contribute not only to precocious OL differentiation but also to the subsequent oligodendroglial loss observed in the 3xTg-AD mice.

### N-MYRF stability is prolonged by Aβ through PKC/GSK3 signaling alterations

To elucidate the mechanism underlying Aβ-induced MYRF accumulation, we first assessed whether Aβ regulates *Myrf* transcription. RT-qPCR analysis of cultured OLs revealed no significant changes in *Myrf* or its upstream regulator *Sox10* mRNA levels following Aβ treatment (**Supplementary Fig. 3.1a**). Consistently, luciferase assays using a construct containing the *Myrf* minimal promoter and SOX10-responsive ECR9 enhancer showed no change in transcriptional activity upon Aβ exposure (**Supplementary Fig. 3.1b**). These results indicate that Aβ does not regulate MYRF expression transcriptionally or via upstream activation, suggesting a post-transcriptional mechanism.

Therefore, we next examined whether Aβ affects MYRF protein stability or degradation. Cycloheximide (CHX) chase assays were performed in OLs treated with vehicle, Aβ, or the GSK3 inhibitor AR-A014418, which is supposed to prevent N-MYRF degradation by blocking GSK3-mediated phosphorylation (Nakayama et al., 2018) (**Fig. 3a**). In control cells, N-MYRF exhibited a half-life of 44.12 ± 10.55 min. As expected, GSK3 inhibition markedly extended MYRF stability (110.08 ± 13.08 min), confirming GSK3’s role in N-MYRF degradation in OLs. Aβ treatment also significantly prolonged N-MYRF half-life (71 ± 9.8 min), indicating that Aβ impairs MYRF degradation (**Fig. 3b, c**).

**Figure 3.**
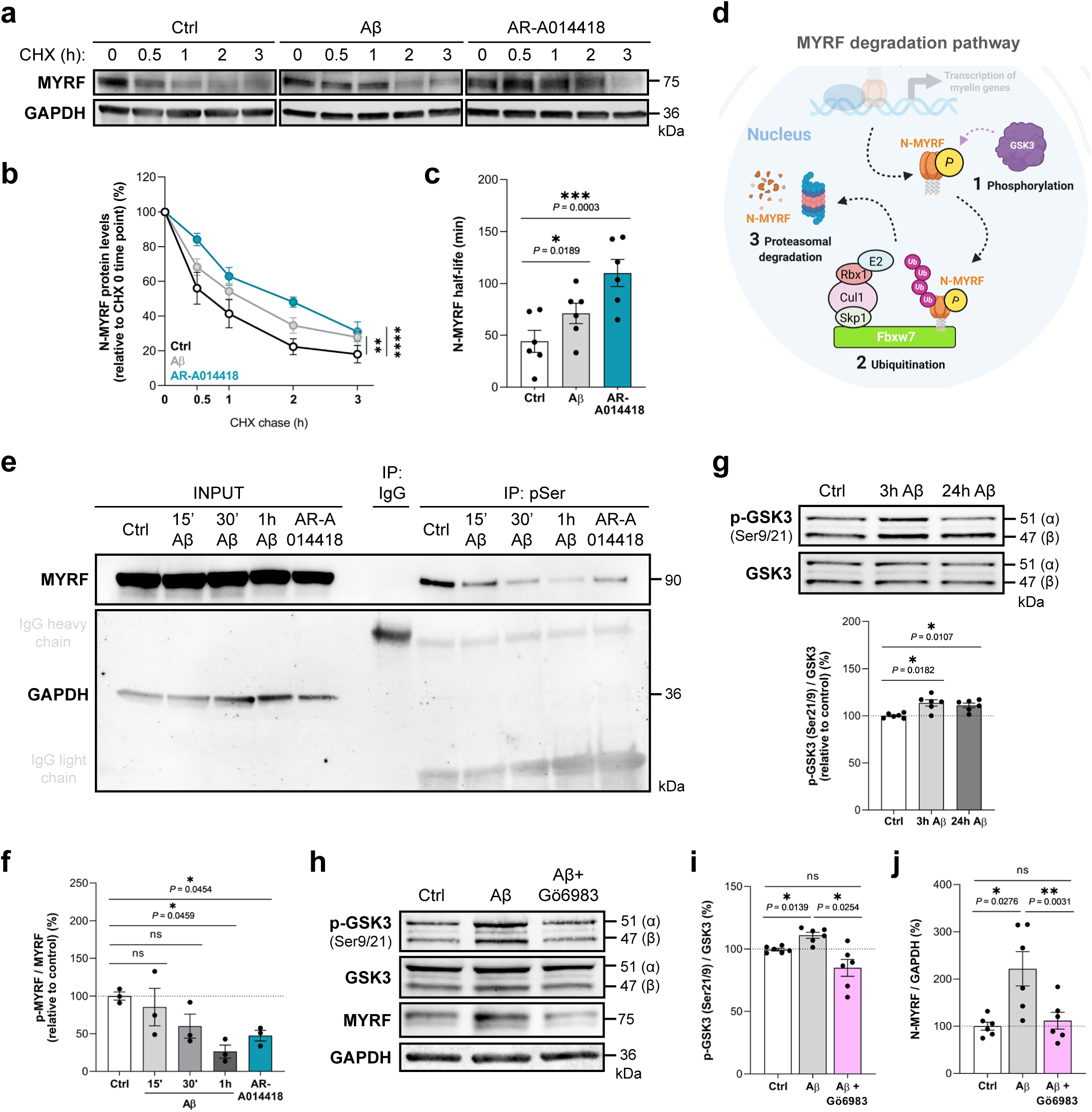
Aβ enhances MYRF protein stability by impairing PKC/GSK3-dependent degradation. (**a-c**) Cycloheximide (CHX) chase analysis of N-MYRF stability in primary OLs treated with vehicle, 1 µM Aβ, or 1 µM AR-A014418 (GSK3 inhibitor; positive control). (**a**) Representative western blots of N-MYRF and GAPDH from OLs incubated with CHX (100 µg/ml) for the indicated times, and (**b**) quantification of N-MYRF levels over time, normalized to the 0 h CHX time point. (**c**) Estimated half-life of N-MYRF in each condition. (**d**) Described degradation pathway of nuclear MYRF: After performing transcriptional activity in the nucleus, N-MYRF is (1) phosphorylated by GSK3, (2) ubiquitinated by the SCF^Fbxw7^ complex, and (3) degraded by the proteasome (adapted from Nakayama et al. 2018; created with BioRender.com). Ub: ubiquitin; P: phosphorylation. (**e-f**) Phosphoserine (pSer) immunoprecipitation (IP) assay of MYRF in HEK293T cells transfected with the MYRF construct and treated with vehicle, Aβ (1 µM; 15 min, 30 min, or 1 h), or AR-A014418 (1 µM). (**e**) Western blot analysis of input and pSer-IP fractions. GAPDH and IgG were used as loading and IP controls, respectively. (**f**) Quantification of pSer-MYRF levels normalized to input and expressed relative to control. (**g**) Western blot analysis of inhibitory Ser21/9 phosphorylation of GSK3 (p-GSK3) in primary OLs treated with vehicle or Aβ for 3 or 24 h, normalized to total GSK3 levels. (**h-j**) Western blot analysis of (**i**) p-GSK3 (normalized to total GSK3) and (**j**) N-MYRF (normalized to GAPDH) in OLs treated with vehicle, Aβ, or Aβ + Gö6983 (PKC pan-inhibitor; 100 nM; added 30 min prior to Aβ) for 24 h. Data are presented as mean ± S.E.M, with dots representing independent experiments. *p<0.05, **p<0.01, ***p<0.001, ****p<0.0001; statistical significance was determined using one- or two-way ANOVA followed by Tukey’s or Dunnett’s *post-hoc* test, as appropriate.

To further dissect the mechanism, we examined the described degradation pathway in which GSK3 phosphorylates N-MYRF at a serine residue, priming it for SCF^Fbxw7^-mediated ubiquitination and proteasomal degradation (Nakayama et al., 2018) (**Fig. 3d**). Whether this pathway operates in OLs, and how Aβ interferes with it, remained unknown. We first assessed MYRF phosphorylation via phosphoserine (pSer) immunoprecipitation in HEK293T cells overexpressing MYRF and treated with Aβ for 15, 30, or 60 min. Interestingly, western blot analysis of pSer-IP fractions revealed a time-dependent decrease in MYRF phosphorylation following Aβ exposure (15 min: 85.46 ± 24.79%; 30 min: 60.13 ± 16%; 1 h: 26.31 ± 8.67%, relative to control) (**Fig. 3e, f**). AR-A014418 treatment similarly reduced pSer-MYRF levels (47.51 ± 7.09%), confirming GSK3 dependence for this phosphorylation. These data suggest that Aβ reduces MYRF phosphorylation, likely via GSK3 inhibition.

GSK3 activity is primarily regulated by phosphorylation: tyrosine phosphorylation activates the kinase (K. Hughes et al., 1993; Lochhead et al., 2006), while inhibitory phosphorylation at Ser21/9 suppresses it (Beurel et al., 2015). Western blot analysis revealed increased Ser21/9 phosphorylation in OLs treated with Aβ for 3 and 24 h (113.59 ± 3.34% and 110.94 ± 2.43%, respectively, relative to control; **Fig. 3g**), consistent with reduced GSK3 activity. To validate this functionally, we measured β-catenin levels, a canonical GSK3 substrate. Aβ treatment significantly increased β-catenin protein levels at 3 h (130.05 ± 8.77% vs 100% control; **Supplementary Fig. 3.2a**), supporting GSK3 inhibition. Moreover, GSK3 activity also depends on proximity to its substrates and subcellular localization (Beurel et al., 2015). Although traditionally considered cytosolic, GSK3 must translocate to the nucleus to phosphorylate some substrates, such as N-MYRF. Immunofluorescence revealed that the nuclear/cytosolic GSK3 ratio was significantly reduced in OLs following Aβ treatment (vehicle: 1.07 ± 0.04 a.u.; Aβ: 0.81 ± 0.02 a.u.; **Supplementary Fig. 3.2b**), indicating diminished nuclear availability of GSK3 to interact with MYRF and thereby limiting its phosphorylation.

Since PKC is a known upstream inhibitor of GSK3 via Ser21/9 phosphorylation (Beurel et al., 2015), and Aβ has been shown to activate PKC in various brain cell types, including neurons (Manterola et al., 2013; Ortiz-Sanz et al., 2022) and OLs (Balantzategi et al., 2025), we next tested whether PKC can modulate Aβ-induced effects in MYRF. PKC inhibition with Gö6983 significantly prevented Aβ-induced GSK3 Ser21/9 phosphorylation (Aβ: 110.94 ± 2.43%; Aβ + Gö6983: 84.91 ± 6.73%; **Fig. 3h, i**), and N-MYRF accumulation (Aβ: 221.64 ± 36.19%; Aβ + Gö6983: 111.64 ± 17.69%; **Fig. 3h, j**), identifying PKC as a key upstream modulator of GSK3-MYRF signaling and highlight Gö6983 as a potential tool to counteract Aβ-induced effects in OLs.

### Aβ promotes Fbxw7-dependent non-proteolytic ubiquitination of MYRF

Building on our findings that Aβ prolongs N-MYRF stability by reducing GSK3-mediated phosphorylation, we next investigated whether Aβ also interferes with the subsequent step of MYRF degradation: ubiquitination by the E3 ligase Fbxw7 (Nakayama et al., 2018). Since phosphorylation is required for Fbxw7 recognition, we asked whether Aβ also alters MYRF ubiquitination.

To address this, HEK293T cells were co-transfected with GFP-MYRF and FLAG-tagged ubiquitin (FLAG-Ub), together with either Fbxw7, Parkin (negative control E3 ligase), or no E3 ligase (ø). Cells were treated with vehicle or Aβ (1 µM, 3 h), and GFP-MYRF was immunoprecipitated using GFPTrap-A beads. MYRF-linked ubiquitin was detected via anti-FLAG immunoblotting (**Fig. 4a**). As expected, Fbxw7 expression was required for robust MYRF ubiquitination, while absence of ligase or Parkin yielded minimal signal (ø: 0.27 ± 0.12 a.u.; Fbxw7: 1.42 ± 0.09 a.u.; Parkin: 0.48 ± 0.23 a.u.; **Fig. 4b, c**). Notably, GFP-MYRF input levels were reduced in Fbxw7-expressing cells, consistent with degradation. Unexpectedly, Aβ treatment significantly increased MYRF ubiquitination in Fbxw7-expressing cells (vehicle: 1.42 ± 0.09 a.u.; Aβ: 2.19 ± 0.15 a.u.; **Fig. 4b, c**), despite also elevating MYRF protein levels (**Supplementary Fig. 4.1a**). This paradox suggests that Aβ promotes ineffective or non-proteolytic ubiquitination of MYRF. Importantly, this Aβ-induced increase was observed only in Fbxw7-expressing cells and exclusively in elution fractions, confirming MYRF- and Fbxw7-specificity.

**Figure 4.**
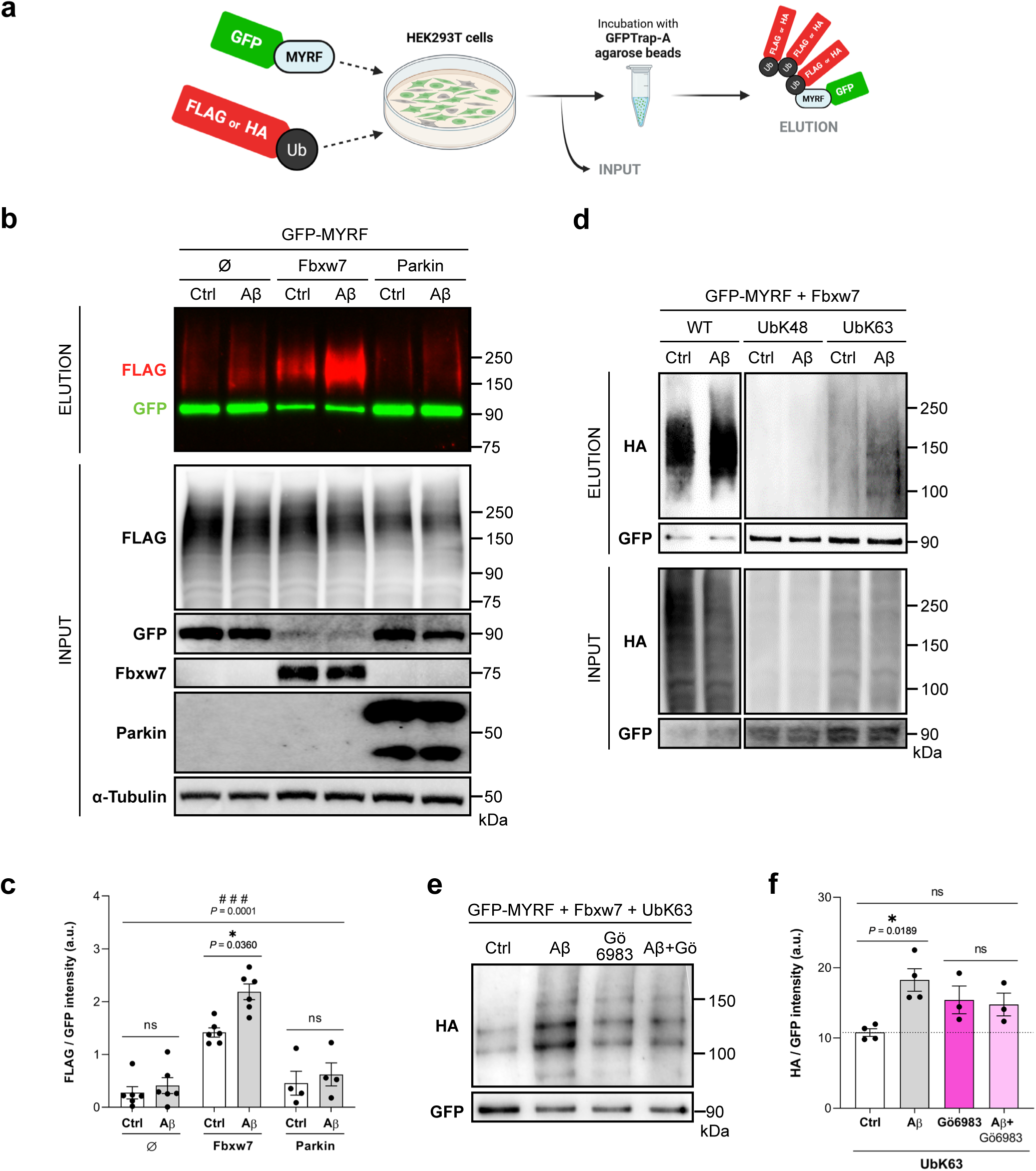
Aβ increases non-proteolytic MYRF ubiquitination in an Fbxw7-dependent manner, an effect prevented by PKC inhibition. (**a**) Schematic representation of the experimental workflow for assessing MYRF ubiquitination: HEK293T cells were co-transfected with GFP-tagged MYRF and either FLAG- or HA-tagged ubiquitin constructs; GFP-MYRF was immunoprecipitated using GFPTrap-A agarose beads; and ubiquitin signal was detected by FLAG or HA immunoblotting. (**b**) Western blot analysis of MYRF ubiquitination after GFP pulldown in cells co-transfected with GFP-MYRF and either no E3 ligase (ø), Fbxw7, or Parkin, and treated with or without Aβ (1 µM, 3 h). Ubiquitinated MYRF was detected with anti-FLAG (red) and total MYRF with anti-GFP (green). Corresponding input blots show expression of FLAG-ubiquitin, GFP-MYRF, Fbxw7, Parkin, and α-Tubulin. (**c**) Quantification of MYRF ubiquitination as FLAG/GFP signal intensity ratio. (**d**) Analysis of Fbxw7-mediated ubiquitin chain specificity using three HA-tagged ubiquitin constructs: WT, UbK48 (favoring proteolytic degradation), and UbK63 (associated with non-proteolytic functions), co-transfected with GFP-MYRF and Fbxw7. Cells were treated with vehicle or Aβ, and ubiquitin signal was detected by anti-HA western blot after GFP pulldown. (**e**) Western blot and quantification (**f**) of non-proteolytic ubiquitination levels of MYRF after GFP pulldown in cells expressing GFP-MYRF, Fbxw7, and HA-UbK63, in the presence or absence of Aβ and/or the PKC inhibitor Gö6983 (100 nM, 30 min pre-treatment). HA-ubiquitin signal was normalized to GFP-MYRF levels. Data are presented as mean ± S.E.M; and dots represent independent experiments. *p<0.05, between treatments; # # #p<0.001, between E3 ligases. Statistical significance was determined by two-way ANOVA followed by Sidak’s *post-hoc* test and one-way ANOVA followed by Tukey’s *post-hoc* test.

To determine the nature of this modification, we next investigated the type of ubiquitin chains involved in MYRF labeling under Aβ exposure. Ubiquitin chains can form through any of its seven lysine residues, resulting in distinct lengths, compositions, and cellular functions. Lysine 48 (K48)-linked chains typically signal proteasomal degradation (Glickman & Ciechanover, 2002; Ihara et al., 2012), whereas K63-linked chains mediate diverse regulatory functions beyond degradation, depending on cellular context and targets (Tramutola et al., 2018). To assess this, we co-transfected GFP-MYRF and Fbxw7 with different HA-tagged ubiquitin constructs: WT-Ub, K48-only Ub (UbK48; favoring proteasomal degradation), or K63-only Ub (UbK63; non-degradative). GFP pulldown assays revealed that Aβ enhanced MYRF-linked ubiquitination with WT-Ub and UbK63, with no changes in UbK48 (**Fig. 4d-f**), indicating a shift toward K63-linked, non-proteolytic ubiquitination, likely contributing to MYRF stabilization. Proteasomal dysfunction was ruled out, since Aβ had no effect on chymotrypsin-like proteasome activity in either HEK293T cells or primary OLs (**Supplementary Fig. 4.1b**), reinforcing the idea that MYRF stabilization results from altered phosphorylation/ubiquitination signaling.

Importantly, this Aβ-induced aberrant ubiquitination was PKC-dependent. Gö6983 pre-treatment prevented Aβ-induced MYRF ubiquitination for both WT-Ub (**Supplementary Fig. 4.1c**) and UbK63 (**Fig. 4e, f**), implicating PKC as a key upstream regulator of this non-canonical ubiquitination pathway.

Interestingly, bulk RNA-seq analysis of MACS-purified O4^+^ (immature and mature) OLs from 6-month-old WT and 3xTg-AD mice also revealed notable deregulation of ubiquitin-dependent (and broader) protein degradation pathways in GO enrichment analysis (**Supplementary Fig. 4.2a-c**), consistent with the altered ubiquitination observed in Aβ-treated cells. Notably, genes such as *Ubb* (ubiquitin precursor) and *Fbxw7*, among others, were upregulated in 3xTg-AD OLs (**Supplementary Fig. 4.2d**), pointing to enhanced ubiquitination machinery. These transcriptomic alterations align with our *in vitro* findings and support the hypothesis that Aβ-induced MYRF accumulation may arise from disrupted degradation signaling.

### PKC inhibition restores oligodendroglial and myelin alterations in the hippocampus of 3xTg-AD mice

Given our molecular evidence that Aβ promotes MYRF accumulation via PKC-dependent inhibition of GSK3 and impaired degradation, we next tested whether pharmacological PKC inhibition could reverse these alterations *in vivo*. To this end, 12—13-month-old WT and 3xTg-AD mice were implanted with osmotic minipumps delivering the broad-spectrum PKC inhibitor Gö6983 (100 nM) or vehicle into the lateral ventricle for 28 days (**Fig. 5a**). Mice were weighed regularly throughout the treatment, and no significant changes in body weight were observed across groups, ruling out adverse systemic effects of the drug (**Supplementary Fig. 5.1a**).

**Figure 5.**
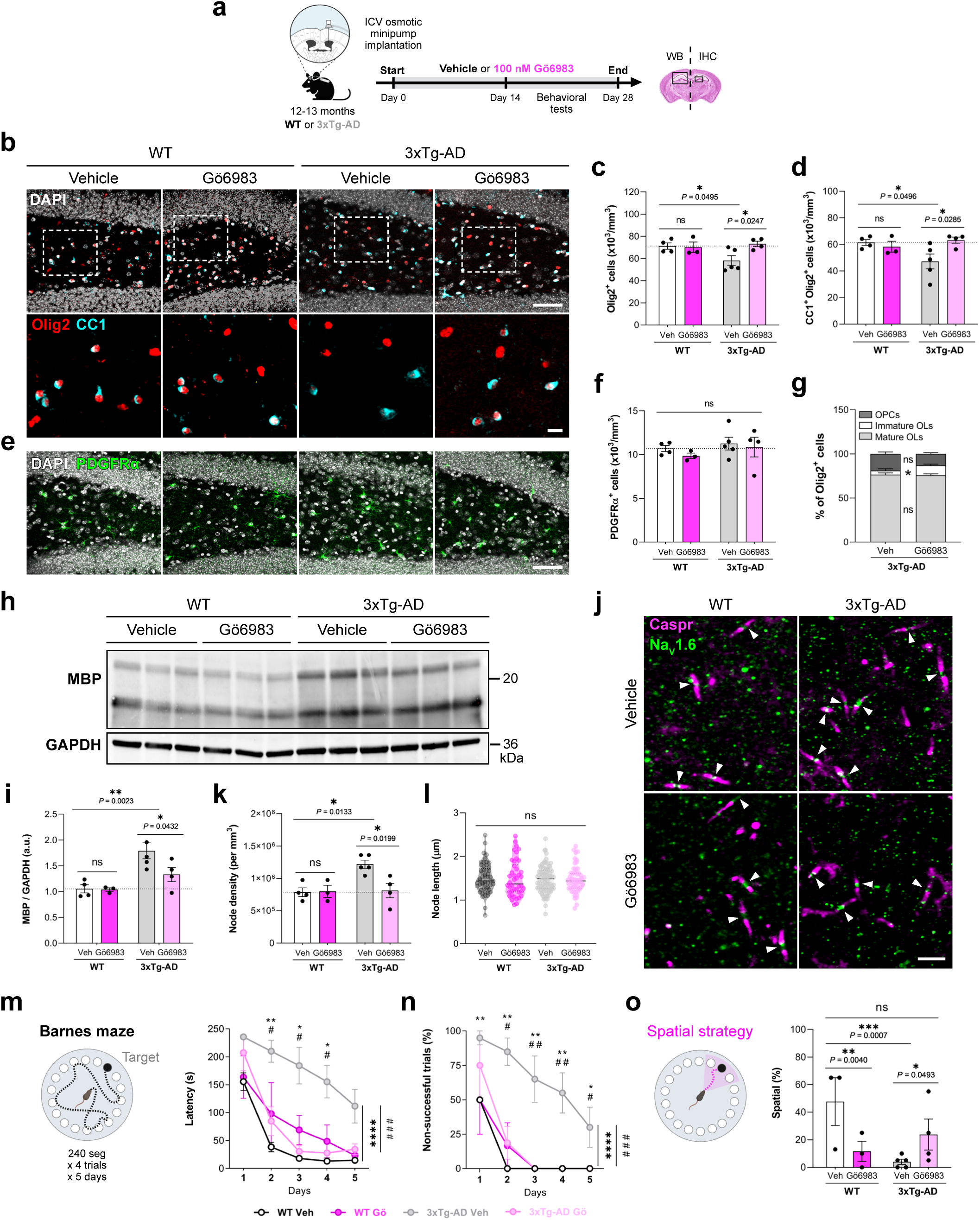
PKC inhibition via intracerebroventricular infusion of Gö6983 restores oligodendroglial populations, myelin integrity, and hippocampal-dependent spatial learning in 3xTg-AD mice. (**a**) Schematic of the experimental design: 12—13-month-old WT and 3xTg-AD mice were implanted with osmotic minipumps delivering Gö6983 (100 nM; 0.11 μl/h) or vehicle into the lateral ventricle for 28 days. Brains were collected for immunohistochemistry (IHC; ipsilateral hemisphere) and western blot (WB; contralateral hemisphere). (**b**) Representative fluorescent images of Olig2 (red), CC1 (cyan), and DAPI (white) in the hilus. (**c**) Quantification of total Olig2^+^ cells and (**d**) mature CC1^+^Olig2^+^ OLs. Scale bars: 50 µm (overview), 10 µm (magnification). (**e**) Representative IHC of PDGFRα^+^ OPCs (green) and DAPI (white); and (**f**) quantification of PDGFRα^+^ OPC density. Scale bar: 50 µm. (**g**) Relative fraction of oligodendroglial subpopulations: PDGFRα⁺ OPCs, immature Olig2⁺CC1⁻PDGFRα⁻ OLs, and mature CC1⁺Olig2⁺ OLs in vehicle- or Gö6983-treated 3xTg-AD mice. (**h**) Representative WB and (**i**) quantification of MBP levels normalized to GAPDH in hippocampal lysates. (**j**) IHC of Caspr (magenta; paranodes) and Na_V_1.6 (green; nodes) in the dentate gyrus; nodes of Ranvier are marked by white arrowheads. (**k**) Quantification of node density and (**l**) node length. Scale bar: 5 µm. (**m**-**o**) Assessment of spatial learning and memory using the Barnes maze: schematic representation and quantitative analysis of (**m**) escape latency, (**n**) percentage of unsuccessful trials, and (**o**) use of spatial search strategy across five test days. Data are presented as mean ± S.E.M; dots represent individual mice (n = 3-6 per group), or individual nodes in the violin plot (**l**). *p<0.05, **p<0.01, ***p<0.001, ****p<0.0001, (**m**, **n**: genotype comparison, WT veh vs 3xTg-AD veh); #p<0.05, # #p<0.01, # # #p<0,001, (**m**, **n**: treatment comparison, 3xTg-AD veh vs 3xTg-AD Gö6983). Statistical significance was determined by two-way ANOVA followed by Sidak’s, Tukey’s, or Bonferroni’s post-hoc tests, as appropriate.

Western blot analysis of contralateral hippocampal lysates confirmed both effective PKC inhibition and adequate diffusion of Gö6983 to both hemispheres, as evidenced by reduced p-PKC/PKC ratio in Gö6983-treated 3xTg-AD mice (vehicle: 5.38 ± 0.72 a.u.; Gö6983: 3.19 ± 0.4 a.u.; **Supplementary Fig. 5.1b, c**). Importantly, this analysis also revealed a significant decrease in MYRF protein levels in Gö6983-treated mice (vehicle: 1.1 ± 0.16 a.u.; Gö6983: 0.85 ± 0.04 a.u.; **Supplementary Fig. 5.1b, d)**, which was corroborated by immunohistochemistry in the ipsilateral dentate gyrus (vehicle: 71.26 ± 2.4%; Gö6983: 62.47 ± 2.56%; **Supplementary Fig. 5.1e, f**), indicating that sustained PKC inhibition effectively counteracts the aberrant MYRF upregulation observed in 12-month-old 3xTg-AD mice.

We next examined whether normalization of MYRF levels by PKC inhibition was accompanied by recovery of oligodendroglial populations. In the dentate gyrus, Gö6983 treatment did not affect oligodendroglial populations in WT mice, but fully restored them to WT values in 3xTg-AD mice. PKC inhibition increased the total number of Olig2^+^ lineage cells (WT veh: 71.25 ± 2.91 x 10^3^ cells/mm^3^; 3xTg-AD veh: 58.18 ± 4.43 x10^3^ cells/mm^3^; 3xTg-AD Gö6983: 73.19 ± 2.42 x10^3^ cells/mm^3^; **Fig. 5b, c**) and mature CC1^+^Olig2^+^ OLs (WT veh: 61.44 ± 2.20 x10^3^ cells/mm^3^; 3xTg-AD veh: 47.14 ± 5.53 x10^3^ cells/mm^3^; 3xTg-AD Gö6983: 63.13 ± 2.31 x10^3^ cells/mm^3^; **Fig. 5b, d**) to WT levels, without affecting PDGFRα^+^ OPC numbers (**Fig. 5e, f**). Notably, the percentage of immature OLs, reduced in 3xTg-AD mice (**Fig. 1j**), was also markedly recovered in 3xTg-AD mice following treatment (**Fig. 5g**). Together, these findings indicate that PKC inhibition prevents OL loss and re-establishes lineage balance.

Given that disrupted OL maturation and survival contribute to altered myelination, we next assessed myelin integrity. Western blot analysis of hippocampal lysates revealed that MBP levels, significantly elevated in 3xTg-AD mice, were reduced toward WT values after Gö6983 treatment (WT veh: 1.05 ± 0.08 a.u.; 3xTg-AD veh: 1.79 ± 0.16 a.u.; 3xTg-AD Gö6983: 1.33 ± 0.14 a.u.; **Fig. 5h, i**). In line with this, immunostaining for Caspr and Na_V_1.6 in the dentate gyrus showed that the increased nodal density observed in 3xTg-AD mice was normalized by Gö6983 (WT veh: 7.87 ± 0.68 x10^5^ nodes/mm^3^; 3xTg-AD veh: 12.20 ± 0.62 x10^5^ nodes/mm^3^; 3xTg-AD Gö6983: 8.12 ± 1.11 x10^5^ nodes/mm^3^; **Fig. 5j, k**), while node length remained unchanged (**Fig. 5l**).

These findings demonstrate that sustained PKC inhibition reduces MYRF accumulation and restores hippocampal oligodendroglial populations, myelin protein expression, and nodal organization in 3xTg-AD mice.

### PKC inhibition improves hippocampal-dependent spatial learning in 3xTg-AD mice

Given the role of OLs and myelin integrity in hippocampal circuit function and cognition (Munyeshyaka & Fields, 2022; Shimizu et al., 2023), we next asked whether Gö6983-induced cellular and molecular recovery translated into behavioral improvement. To this end, WT and 3xTg-AD mice treated intracerebroventricularly with Gö6983 or vehicle were subjected to a battery of behavioral tests (**Fig. 5a** and **Supplementary Fig. 5.2a**).

The most prominent effect was observed in the Barnes maze (BM), a hippocampal-dependent task that requires the use of spatial navigation strategies (Harrison et al., 2006). In this test, mice learn the location of a hidden escape box using distal visual cues, with performance assessed over five consecutive days (**Fig. 5m**). Vehicle-treated 3xTg-AD mice showed impaired spatial learning, evidenced by longer escape latencies, higher proportions of unsuccessful trials (**Fig. 5n**), and markedly reduced reliance on hippocampal-dependent spatial strategies (WT veh: 47.67 ± 17.33%; 3xTg-AD veh: 4 ± 1.87%; **Fig. 5o**). Remarkably, Gö6983 treatment significantly improved these parameters: 3xTg-AD mice receiving the PKC inhibitor displayed progressive learning across sessions, reaching WT-like performance in latency (**Fig. 5m**) and success rate (**Fig. 5n**). Although Gö6983 reduced spatial strategy use in WT mice (11.67 ± 7.27%), it partially restored spatial strategy use towards control levels in 3xTg-AD mice (23.75 ± 11.25%; **Fig. 5o**), thereby improving hippocampal-dependent performance. Importantly, this cognitive rescue occurred without altering innate locomotor activity or anxiety-like behavior in either genotype, as assessed in the open field (**Supplementary Fig. 5.2b-d**), elevated plus maze (**Supplementary Fig. 5.2e-g**), and light/dark box (**Supplementary Fig. 5.2h**) paradigms. Thus, the improvement in BM performance cannot be attributed to changes in general activity or emotional state.

Finally, emotional learning and memory were evaluated using the contextual fear conditioning paradigm, and although 3xTg-AD mice exhibited impaired contextual and cued fear memory relative to WT controls, Gö6983 treatment did not significantly reverse these deficits (**Supplementary Fig. 5.2 i**).

Together, these results indicate that PKC inhibition selectively restores hippocampal-dependent spatial learning and memory in 3xTg-AD mice, while leaving anxiety-like behavior and amygdala-dependent emotional memory largely unaffected.

## DISCUSSION

AD has traditionally been defined as a disorder of neuronal loss and synaptic dysfunction, yet converging evidence indicates that OLs and myelin integrity contribute meaningfully to disease progression (Kedia & Simons, 2025). White matter abnormalities have been reported in patients and models, sometimes preceding overt neuronal degeneration, and several studies point to glial dysfunction as an early feature of the disease. Within this broader framework, our data support a model in which Aβ exposure is associated with MYRF dysregulation and altered OL lineage balance and myelination.

In 12-month-old 3xTg-AD mice, the dentate gyrus shows reduced total Olig2^+^ and mature OLs, alongside an increased proportion of newly matured OLs among the EdU-labeled lineage. Observations of pyknotic nuclei and occasional microglial engulfment are consistent with OL loss. These findings align with human *post-mortem* and murine work reporting decreased OL lineage cells in hippocampus (Behrendt et al., 2013; DeFlitch et al., 2022). At the same time, other AD models such as APP/PS1 mouse show increased oligodendrogenesis (Ferreira et al., 2020) and new myelin formation (Chen et al., 2021), particularly at early stages. Taken together, these datasets suggest that although OL differentiation is stimulated, lineage balance is not maintained. We hypothesize that mature OLs undergo cell death, and that the compensatory drive to replace them through accelerated maturation gradually depletes intermediate subpopulations, as observed in our 3xTg-AD mice. Thus, despite OPCs apparently retaining their potential to proliferate and differentiate, turnover and loss outpace renewal. In this scenario, oligodendrogenesis becomes a compensatory but ultimately insufficient response, leading over time to reduced density of myelinating OLs and increased vulnerability of hippocampal circuits.

In this context, MYRF—a transcription factor essential for terminal OL differentiation and myelination—emerges as a key factor. We found that Aβ increases MYRF protein abundance both *in vitro* and *in vivo* and, in cultured OLs, augments its transcriptional activity, resulting in increased MBP levels. Importantly, this observation is consistent with human data from the SEA-AD atlas, which reported a transient upregulation of MYRF early in disease (Gabitto et al., 2024), and with neuropathological studies showing increased *MYRF* mRNA expression in the white matter of AD patients (Ferrer & Andrés-Benito, 2020). While MYRF upregulation may initially seem advantageous for the cell due to its remyelinating potential, our *in vitro* data indicate that sustained elevation reduces OL viability, suggesting that prolonged nuclear stabilization of MYRF may become toxic for the cell, potentially contributing to lineage imbalance and OL loss observed in 3xTg-AD mice. This notion is reinforced by the specific expression of MYRF during differentiation and myelination stages, precisely the subpopulations we found to be more vulnerable to AD-related disruptions compared to OPCs. It is conceivable that Aβ-induced sustained MYRF expression may alter its transcriptional specificity engaging non-physiological or nonspecific target genes, potentially engaging pathways that compromise OL survival. Nevertheless, this hypothesis requires further investigation.

The literature on Aβ toxicity to OLs has reported variable results, likely reflecting differences in dose, exposure time, oligomer species, and cellular context. Increased pyknotic nuclei have been described after short, higher-dose exposures (Desai et al., 2010), selective vulnerability of mature OLs under prolonged soluble Aβ (Horiuchi et al., 2012), reduced MBP^+^ cells at extended treatments (Zhang et al., 2018), and enhanced viability, differentiation and increased MBP expression at lower doses (Gaminde-Blasco et al., 2025; Quintela-López et al., 2019)(Gaminde-Blasco et al., 2025)(Gaminde-Blasco et al., 2025)(Gaminde-Blasco et al., 2025). Our data are consistent with a biphasic interpretation: modest, transient Aβ increases MYRF activity and OL differentiation, whereas persistent elevation and stabilization of MYRF under Aβ-linked signaling may become harmful. This framework does not resolve all contradictions but provides a coherent lens to interpret context-dependent outcomes.

MYRF and OL alterations observed in the 3xTg-AD mice should also be considered within the broader framework of adaptive versus maladaptive myelination. The precise and coordinated production of myelin is essential for the correct functioning of the CNS. Adult-born OLs can support activity-dependent plasticity by myelinating previously unmyelinated axons, extending existing sheaths, and remodeling nodal organization (E. G. Hughes et al., 2018). However, under certain conditions, premature or excessive OL differentiation may disrupt myelin homeostasis, reduce conduction speed, and impair circuit function, a pattern linked to cognitive decline during aging (Bowley et al., 2010; Lasiene et al., 2009; Young et al., 2013). Therefore, induction of MYRF expression by Aβ may lead to hypermyelination, but this synthesized myelin may not be physiological or functionally normal. Consistent with this, we observe increased MBP protein levels and nodal density in 3xTg-AD hippocampus, accompanied by poorer Barnes maze performance; after intervention with Gö6983, normalization of myelin parameters was associated with improved spatial learning. These associations support the concept that excess or aberrant myelin remodeling can be maladaptive in AD contexts. Indeed, defective myelin sheaths have been proposed to be more detrimental to axonal function than their absence, as aberrant myelin uncouples OL support for axons (Schäffner et al., 2023; Simons & Nave, 2016). Myelin deficits have also been linked to enhanced Aβ deposition, exacerbating AD pathology (Depp et al., 2023).

Mechanistically, our results support an altered PKC/GSK3/Fbxw7/MYRF pathway in which Aβ exposure is associated with increased inhibitory Ser21/9 phosphorylation of GSK3, reduced phosphoserine labeling of MYRF, and prolonged stability of the nuclear MYRF fragment. Under our experimental conditions, Aβ appears to reduce GSK3 activation, which contrasts with most reports describing GSK3 overactivation in AD (Lauretti et al., 2020; Zhao et al., 2024); this may again reflect the biphasic nature of Aβ effects and AD. In fact, GSK3β inhibition has been shown to promote OL differentiation via the canonical Wnt/β-catenin pathway (Azim & Butt, 2011), consistent with our findings. We also describe an Fbxw7-dependent shift toward K63-linked (non-proteolytic) ubiquitination of MYRF under Aβ exposure, consistent with stabilization rather than proteasomal degradation. A recent research work reported that the E3 ubiquitin ligase RNF220 targets Olig1 and Olig2 for K63 ubiquitination and stabilization during oligodendroglial development (Y. Li et al., 2024), underscoring the importance of K63 ubiquitination in OL biology. Overall, we propose that Aβ interferes with PKC/GSK3 signaling and Fbxw7-mediated ubiquitination, leading to ineffective clearance of MYRF and aberrantly increased transcriptional activity that disrupts OLs and myelin.

Our results build on prior zebrafish work where PKC modulation impacted OL differentiation and myelination (Balantzategi et al., 2025); here, we extend those findings by delineating a mechanistic cascade in mammalian OLs and by demonstrating associated improvements in cognition. Importantly, sustained PKC inhibition in 3xTg-AD mice reduced MYRF levels in hippocampus, restored OL populations and intermediate subpopulations, normalized myelin protein expression and nodal density, and improved hippocampal-dependent spatial learning, without affecting locomotion or anxiety-like behavior. Given that PKC is a master regulator of numerous signaling pathways, we view Gö6983 as a tool for pathway dissection rather than a therapeutic candidate per se. Nonetheless, the rescue supports the tractability of this axis and motivates investigation of more selective strategies that modulate the PKC/GSK3/Fbxw7/MYRF pathway. As a next step, cell-type-targeted modulation *in vivo* prioritizing OLs but also considering neurons and other glia could clarify the relative contributions of each cell type and help refine therapeutic windows. We acknowledge that our study was conducted in murine models, and validation in human tissues and longitudinal cohorts will be essential to strengthen translational relevance. However, the report of an early MYRF increase followed by later decline in human AD (Gabitto et al., 2024) is consistent with our observations and hypothesis, and we believe that provides an encouraging anchor point. In addition, this study opens an exploratory field into the role of MYRF, and its dysregulation, across other neurodegenerative disorders.

In conclusion, our findings support the view that OLs are active participants in AD and that MYRF dysregulation provides a key connection between Aβ, OL lineage dynamics, myelin remodeling, and cognition. By delineating a PKC-dependent pathway that converges on GSK3-regulated phosphorylation and Fbxw7-mediated ubiquitination of MYRF, we offer a mechanistic framework compatible with early compensatory differentiation and later vulnerability. The fact that modulating this axis restores cellular and behavioral readouts in 3xTg-AD mice suggests that preserving white matter integrity, alongside neuronal protection, could be integral to maintaining cognitive function in AD.

## Acknowledgements

We thank L. Escobar, L. Cortés, V. Montoya and Z. Martínez for technical support. We are grateful to A. Aransay from the CIC BioGUNE Genome Analysis Platform for assistance with RNA library preparation and sequencing. We also thank B. Emery (Jungers Center for Neurosciences Research, OHSU, Portland, USA) for providing the double-tagged MYRF plasmid and *Mbp* and *Mag* luciferase reporters, and for valuable experimental advice; Y. Park (Jacobs School of Medicine, UB, Buffalo, USA) for the GFP-MYRF construct; and M. Wegner (Institut für Biochemie, FAU, Erlangen, Germany) for the *Myrf* promoter luciferase plasmid. Graphical schematics were created with BioRender.com.

## Author contributions

U B designed and performed the experiments, analyzed and interpreted data, and wrote the manuscript. A G-B, N A, J R, R S-G, N H-P, M F-B, F N S and E C-Z performed the experiments, analyzed and interpreted data. A G F, F G-M, U M, E S-G and JL Z provided advice on experimental design and data interpretation. E A contributed to the conception and design of the project, analyzed and interpreted data and reviewed the manuscript. All authors have read and approved the final manuscript.

