## Supplementary Material for "PKC-dependent MYRF dysregulation links Aβ pathology to oligodendrocyte, myelin and cognitive alterations in Alzheimer’s disease"

### SUPPLEMENTARY METHODS

#### Immunofluorescence

*Immunohistochemistry of mouse brain tissue*

IHC was performed following the protocol described in the *Materials and Methods* section, using primary antibodies against Olig2 (ms $IgG_{2a}$, 1:500; #MABN50, Millipore) and Iba1 (gp, 1:500; #234308, Synaptic Systems). Images were acquired using a Leica TCS STED CW SP8X confocal microscope using a 63× oil-immersion objective.

*Immunocytochemistry of HEK293T cells*

HEK293T cells were fixed in 4% PFA for 10 min at RT, and ICC was performed following the same protocol described for primary OLs. Primary antibodies included anti-Myc tag (ms $IgG_1$, 1:500; #05-724, Millipore) and anti-FLAG tag (rb, 1:1,000; #2368, Cell Signaling).

*Immunocytochemistry of primary oligodendrocytes*

OLs were exposed to vehicle or Aβ when indicated. ICC was performed following the same protocol described in the *Materials and Methods* section, using the following primary antibodies: anti-MBP (ch, 1:200; #AB9348, Millipore), anti-MYRF (rb, 1:100; #ABN45, Millipore), and anti-GSK3α/β (ms $IgG_{2A}$, 1:500; #sc-7291, Santa Cruz).

#### Western blot

Primary OL and mouse hippocampi samples were prepared as described in the *Materials and Methods* section. For HEK293T cells, 48 h post-transfection and after treatments, cells (seeded at 5 x$10^4$ - 1 x$10^5$ per well) were gently scraped in their culture medium and transferred into clean tubes. Samples were centrifuged (500 x g, 6 min, 4 °C), pellets rinsed with PBS, and centrifuged again under the same conditions to remove residual serum. The resulting pellets were resuspended in RIPA lysis buffer supplemented with protease and phosphatase inhibitors (ThermoFisher), incubated on ice for 30 min, and centrifuged (12,000 x g, 10 min, at 4 °C). Supernatants were carefully collected, mixed with 4x SDS sample buffer, and boiled at 95 °C for 5 min.

Western blotting was performed as described in the *Materials and Methods* section, using primary antibodies against Myc tag (ms, 1:2,000; #05-724, Millipore), FLAG tag (rb, 1:1,000; #2368, Cell Signaling), MBP (ms, 1:1,000; #SMI-99, BioLegend), β-catenin (rb, 1:1,000; #9562, Cell Signaling), p-PKC (rb, 1:1,000; #9371, Cell Signaling), PKC (rb,

1:1,000; #ab179521, abcam), MYRF (rb, 1:1,000; #ABN45, Millipore), and GAPDH (ms, 1:5,000; #MAB374, Millipore).

**Quantitative Real time-Polymerase Chain Reaction (RT-qPCR)**

Total RNA was extracted from cultured primary oligodendrocytes using the RNA Mini Kit (Qiagen) following the manufacturer's protocol. RNA concentration and integrity were assessed with a NanoDrop™ 2000 spectrophotometer (Thermo Fisher Scientific). Complementary DNA was synthesized using Superscript™ III reverse transcriptase (Invitrogen) and random primers in a Verity Thermal Cycler (Applied Biosystems). Specific primers for *Myrf* (forward: 5'-GTGGGCGACCCTATATGATG-3'; reverse: 5'-TCCACACGATAGGTGAGCAT-3') and *Sox10* (forward: 5'-ATGTCAGATGGGAACCCAGA-3'; reverse: 5'-GTCTTTGGGGTGGTTGGAG-3') were newly designed and synthesized by Integrated DNA Technologies (IDT). RT-qPCR reactions were run in triplicate with 5 µl SYBR Green Master Mix (Bio-Rad), 0.5 µl cDNA, 1 µl primer mix, and 3.5 µl RNase-free water. Amplification was performed on a CFX96 Touch Real-Time PCR Detection System (Bio-Rad) under the following program: 95 °C for 3 min, then 40 cycles of 95 °C for 10 s and 60 °C for 30 s. Product specificity was verified by melting curve analysis. Data were normalized using geNorm software based on the expression of three housekeeping genes (*Gapdh*, *B2m*, and *Pgk1*).

**Dual-luciferase assays**

Luciferase assay using the pGL4.10_myrfprom_ECR9 reporter construct was performed in vehicle- or Aβ-treated primary OLs, following the protocol described in the *Materials and Methods* section. *Firefly* luciferase activity driven by the ECR9 regulatory element was normalized to *Renilla* activity, and values were expressed relative to empty vector.

**Chymotrypsin-like proteasome activity assay**

HEK293T cells and primary cultured oligodendrocytes were seeded in 96-well plates at $1x10^4$ cells per well and treated with vehicle or Aβ (1 µM) for 3 or 24 h, with or without MG132 (1 µM, 1 h) as a positive control for proteasome inhibition. Proteasome activity was measured using the Proteasome-Glo™ Cell-Based Assay (Promega) according to the manufacturer's protocol. Briefly, Proteasome-Glo™ Cell-Based Reagent was added to each well at a 1:1 ratio, incubated with gentle shaking for 2 min, and further incubated for 10 min at RT. Luminescence was measured using the CLARIOstar® Plus microplate reader (BMG Labtech). Data were normalized to vehicle-treated cells (set as 100%).

**Magnetic activated cell sorting (MACS)**

For transcriptomic analysis of oligodendrocytes in early Alzheimer's disease, $O4^+$ oligodendrocytes were isolated from 6-month-old WT and 3xTg-AD mice. Two animals were pooled per sample ($n^{WT}$ = 4 (8 animals), $n^{3xTg\text{-}AD}$ = 3 (6 animals)). Mice were anesthetized with isoflurane and euthanized by decapitation, and oligodendrocyte isolation was carried out following previously established protocols (de la Fuente et al., 2020). Brain tissue was dissected, placed in Hibernate A medium (Gibco), and cut into ~1 $mm^3$ pieces. Samples were enzymatically digested with papain (33 U/ml) and DNase I (0.04 mg/ml) in Hibernate A for 30 min at 37 °C, followed by washing with HBSS-/- (Gibco). Tissue was mechanically dissociated in Hibernate A supplemented with B27 (Gibco) and sodium pyruvate (2 mM; Gibco), using a polished glass Pasteur pipette. The resulting cell suspension was filtered through a 70 µm strainer (VWR), and centrifuged at 800 × g for 20 min in 22.5% Percoll (GE Healthcare) with DMEM Glutamax (Gibco). After centrifugation, the Percoll solution was aspirated, and cells were washed with HBSS-/- (Gibco). Then, cells were incubated with anti-O4 antibody (Miltenyi Biotec) for 15 min on ice, washed again with HBSS-/- to remove excess antibody, and magnetically sorted using MS columns (Miltenyi Biotec) according to the manufacturer's instructions. Sorted $O4^+$ cells were eluted in $SATO^+$ medium and processed for RNA extraction and sequencing.

**Bulk RNA sequencing**

RNA was extracted from MACS-isolated $O4^+$ oligodendrocytes using the RNA Micro Kit (Qiagen). Quantity and integrity were assessed with the Qubit RNA Assay Kit (Invitrogen) and the Agilent 2100 Bioanalyzer (Agilent RNA 6000 Pico chips). Libraries were prepared with the NEBNext Single Cell/Low Input RNA Library Prep Kit for Illumina (#E6420S, New England Biolabs) and NEBNext Multiplex Oligos for Illumina (Index Primers 1-12) (#E7335S, New England Biolabs), following the corresponding instruction manual (Version5.0_5/20). The protocol started with 10 ng total RNA when available (otherwise, entire sample). Full-length cDNA was generated by template switching, followed by cDNA amplification and cleanup; 1 µl amplified cDNA was evaluated on an Agilent High Sensitivity DNA chip (#5067-4626, Agilent Technologies). Fragmentation, end repair, tailing, Illumina adapter ligation, and barcoding PCR were performed per kit instructions, with amplification cycles adjusted to input cDNA amount. Libraries were visualized on the Agilent 2100 Bioanalyzer and quantified with the Qubit dsDNA HS DNA Kit (#Q32854, Thermo Fisher Scientific). Base calls (BCL) were converted into FASTQ files utilizing Illumina Inc.'s package bcl2fastq, and quality control was performed with FastQC (Andrews, 2020). Reads were aligned with STAR v2.7.1 (Dobin et al., 2013) against the Ensembl genome of *Mus musculus* (GRCm39.dna.primary_assembly.fa and GRCm39.104.gtf), and expression counts were obtained using htseq-count (-s no) (Anders et al., 2015). The count matrix was imported to R v4.2.2, where low count genes (< 2) were filtered out and expression levels were normalized and further analyzed with DESeq2 (Love et al., 2014). To obtain differentially expressed genes, the contrast "Genotype_AD_vs_WT" was employed in the linear model ~ Day + Genotype, which also accounted for batch effects ("Day"). For data visualization and functional enrichment, *ggplot2* (Wickham, 2016), clusterProfiler (Wu et al., 2021; Yu et al., 2012), and *ggVennDiagram* (Gao et al., 2021) were employed. The code used is available in https://github.com/rodrisenovilla/Balantzategi2026, and raw data is stored in the ENA repository under project PRJEB105099.

**Behavioral tests**

Mice carrying osmotic minipumps were subjected to a battery of behavioral tests during the last 14 days of treatment. All tests were conducted during the light phase, and animals were acclimated to the testing room for 20–30 min prior to each session. All trials were recorded and analyzed using ANY-maze software (ANY-maze; Dublin, Ireland).

*Open Field Test (OF)*

Exploratory activity and anxiety-like behavior were assessed in a square arena (40 x 40 cm, with transparent walls 40 cm high) during a single 5-min trial. Mice were placed in the center and allowed to explore freely. Distance traveled, immobility time, and time spent in the center and periphery were recorded using ARDUCam iDS and ANY-maze software. Anxiety index (AI) was calculated as:

$$AI = \frac{((time\ spent\ in\ the\ safe\ zone\colon periphery) - (time\ spent\ in\ the\ anxiogenic\ zone\colon center))}{((time\ spent\ in\ the\ safe\ zone\colon periphery) + (time\ spent\ in\ the\ anxiogenic\ zone\colon center))}$$

*Elevated Plus Maze (EPM)*

To evaluate anxiety-like behavior, mice were individually place in the center of a four-arm (30 × 5 cm) construction in which two facing arms are open, whereas the other two are enclosed by 30-cm-high walls, for 5 min. The entire plus maze is elevated on a pedestal. Distance traveled, immobility time, and time spent in open (OA) and closed (CA) arms were recorded using ANY-Maze. AI was calculated as:

$$AI = \frac{((time\ in\ CA) - (time\ in\ OA))}{((time\ in\ CA) + (time\ in\ OA))}$$

*Light and dark box (LDB)*

Anxiety-like behavior was assessed in a two-compartment box: an open from above (light) and a closed from every side (dark) one connected to each other by a small gate. Mice were individually placed in the light zone and allowed to freely explore both compartments for 5 min. Time spent in each zone was recorded using ANY-maze. AI was calculated as:

$$AI = \frac{((time\ in\ dark\ box) - (time\ in\ light\ box))}{((time\ in\ dark\ box) + (time\ in\ light\ box))}$$

*Fear conditioning (FC)*

Emotional memory was assessed using a classical conditioning paradigm. This task relies on the ability of animals to associate a neutral stimulus (tone) with an aversive stimulus (mild footshock) (Rustay et al., 2008). A 3 day protocol was employed. On the acquisition day (ACQ), mice were individually placed in the conditioning chamber and allowed to habituate for 1 min to the environment and background white noise. They then received three tone–shock pairings, each consisting of a 10-s auditory cue immediately followed by a 2-s footshock (0.7 mA), separated by 20-s intervals. The session ended with 1 min of white noise. At 24 h (T1) and 48 h (T2) post-training, mice were re-exposed to the conditioned tone without footshock in a modified context to dissociate associative memory from contextual cues. Each test session lasted 5 min: 1 min of white noise, 3 min of continuous tone, and 1 min of white noise. Freezing behavior was recorded using POLY software.

### SUPPLEMENTARY RESULTS AND FIGURES

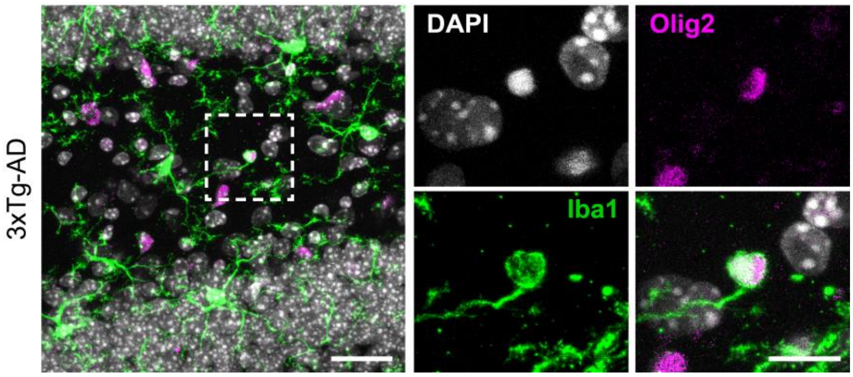


**Supplementary Fig. 1.** ***The 3xTg-AD dentate gyrus exhibit occasional microglial phagocytosis of oligodendrocytes.*** Representative confocal image from a 3xTg-AD mouse showing a microglial cell (Iba1, green) engulfing an Olig2$^{+}$ oligodendrocyte (magenta) with a pyknotic nucleus (DAPI, white) in the dentate gyrus. Scale bars: 25 µm (overview) and 10 µm (zoomed insets).

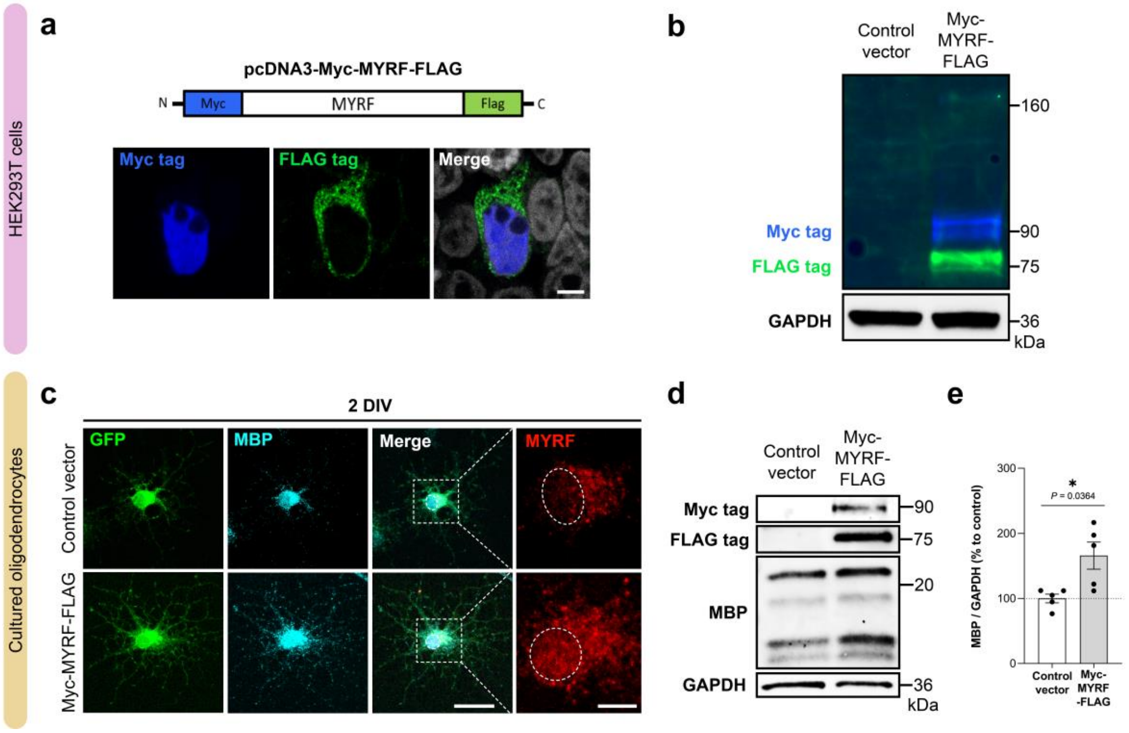


**Supplementary Fig. 2.** ***In vitro characterization of MYRF using the Myc-MYRF-Flag construct.*** (**a**) Schematic of the dual-tagged MYRF expression vector (pcDNA3-Myc-MYRF-FLAG), containing a Myc tag at the N-terminus and a FLAG tag at the C-terminus. HEK293T cells transfected with the construct were stained with anti-Myc tag (blue) and anti-FLAG (green) antibodies to visualize the N-terminal fragment (N-MYRF; nuclear localization) and C-terminal fragment (C-MYRF; ER localization), respectively. Nuclei were stained with DAPI (gray). Scale bar: 5 µm (**b**) Representative fluorescent western blot showing full-length MYRF (fl-MYRF; ~160 kDa) and its cleavage products: N-MYRF (~90 kDa; Myc tag, blue) and C-MYRF (~75 kDa; FLAG tag, green) in HEK293T cells. (**c**) Immunofluorescence images of primary cultured rat oligodendrocytes co-transfected at 2DIV with EGFP and either control empty vector or Myc-MYRF-FLAG, stained for MBP (cyan), and MYRF (red); GFP (green) was also imaged. White dotted lines outline nuclei. Scale bars: 20 µm and 5 µm. (**d**) Western blot analysis of Myc, FLAG, MBP, and GAPDH, and (**e**) quantification of MBP expression in 2 DIV oligodendrocytes transfected with control or Myc-MYRF-FLAG plasmid. MBP levels were normalized to GAPDH and expressed as percentage relative to control. Data are shown as mean ± S.E.M, with dots representing individual experiments. *$p<0.05$; statistical significance was determined by two-tailed paired t-test.

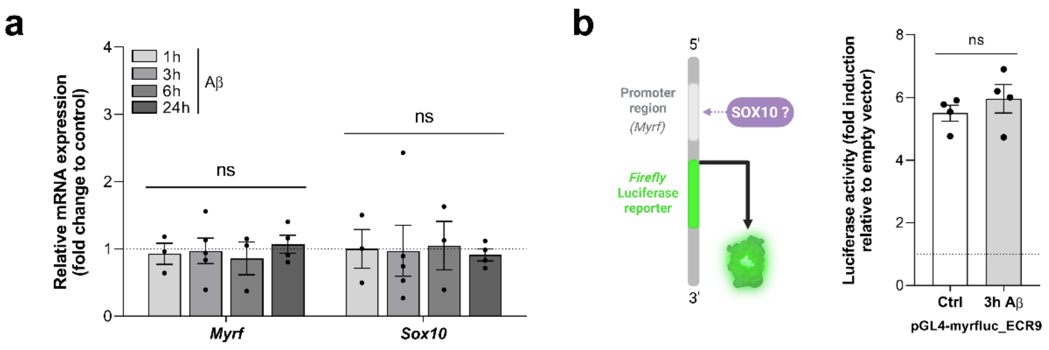


**Supplementary Fig. 3.1.** ***Aβ does not alter MYRF transcription.*** (**a**) RT-qPCR analysis of *Myrf* and *Sox10* mRNA levels in primary cultured OLs at 3 DIV, treated with vehicle or 1 µM Aβ for 1, 3, 6, or 24 h. (**b**) Luciferase reporter assay using the *myrfluc_ECR9* construct (containing a minimal *Myrf* promoter) in 3 DIV OLs treated with or without Aβ for 3 h. Luciferase activity is expressed as fold induction relative to empty vector. Data are presented as mean ± S.E.M., with dots representing independent experiments. Statistical significance was assessed using two-tailed paired t-tests (**b**) or one-way ANOVA followed by Dunnett's *post-hoc* test (**a**).

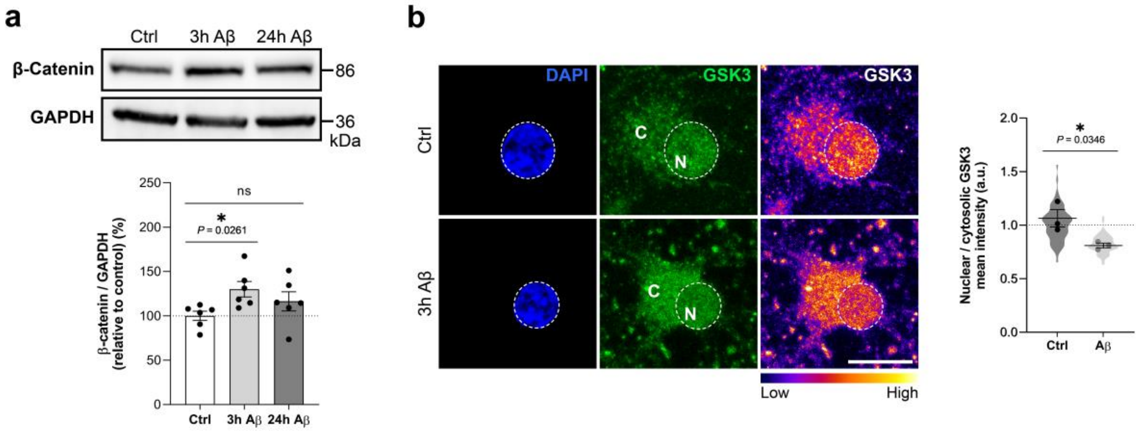


**Supplementary Fig. 3.2.** ***Aβ alters GSK3 activation and subcellular localization in cultured oligodendrocytes.*** (**a**) Western blot analysis of β-catenin levels in primary oligodendrocytes treated with vehicle or 1 µM Aβ for 3 or 24 h, normalized to GAPDH and expressed as percentage relative to control. (**b**) Quantification of the nuclear-to-cytosolic (N/C) GSK3 ratio by immunofluorescence in vehicle- and Aβ-treated OLs. Fluorescence images show GSK3 expression (green and fire LUT) and nuclei stained with DAPI (blue; outlined with white dashed circles). Intensity scale bar corresponds to the fire LUT of GSK3. Scale bar: 10 µm. Data are presented as mean ± S.E.M., with dots representing independent experiments; violin plot reflects quantification of individual cells. *$p<0.05$; statistical significance was assessed using two-tailed nested t-test (**b**) or one-way ANOVA followed by Dunnett's *post-hoc* test (**a**).

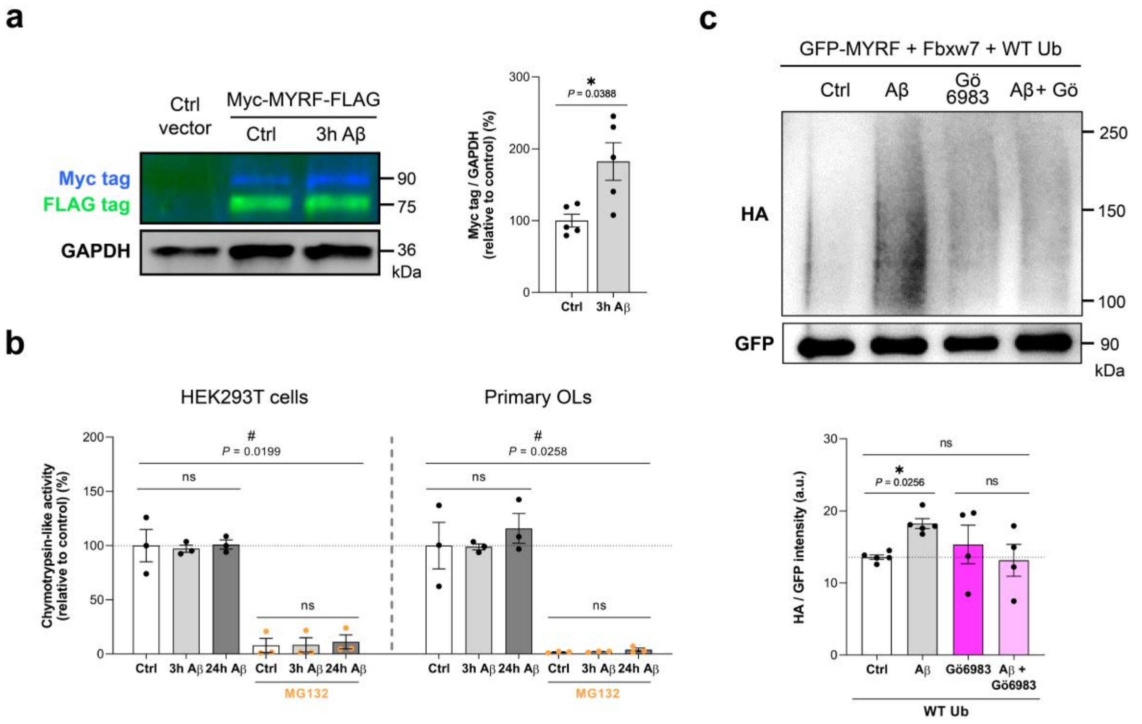


**Supplementary Fig. 4.1.** (**a**) Western blot analysis and quantification of N-MYRF levels in HEK293T cells transfected with either control vector or Myc-MYRF-FLAG construct, and treated with vehicle or Aβ (1 µM, 3 h). N-MYRF was detected using an anti-Myc tag antibody (blue), and C-MYRF with anti-FLAG tag antibody (green).Protein levels were normalized to GAPDH and expressed as percentage relative to control. (**b**) Chymotrypsin-like proteasome activity in HEK293T cells and primary OLs treated with vehicle or Aβ for 3 or 24 h. The canonical proteasome inhibitor MG132 (1 µM, 1 h) was used as a positive control, validating the assay. (**c**) Quantification of MYRF ubiquitination levels by GFP pulldown in HEK293T cells co-expressing GFP-MYRF, Fbxw7, and HA-tagged WT ubiquitin, treated with vehicle, Aβ, Gö6983 (100 nM), or Aβ + Gö6983. HA-ubiquitin signal was normalized to GFP-MYRF levels. Data are presented as mean ± S.E.M, with dots representing individual experiments. *p<0.05, #p<0.05; statistical significance was determined using two tailed paired t-test, one-way ANOVA followed by Tukey's *post-hoc* test, or two-way ANOVA followed by Sidak's or Dunnett's *post-hoc* test, as appropriate.

**Transcriptomic analysis of MACS-isolated O4+ oligodendrocytes from 6-month-old WT and 3xTg-AD mice**

Bulk RNA-seq analysis of MACS-purified $O4^+$ oligodendrocytes from 6-month-old WT and 3xTg-AD mice revealed extensive transcriptional alterations, including dysregulation of protein degradation pathways (**Supplementary Fig. 4.2a**). A total of 5,537 differentially expressed genes (DEGs) were identified, with 2,392 upregulated and 3,145 downregulated in 3xTg-AD OLs (DeSeq2, adjusted $P<0.05$) (**Supplementary Fig. 4.2b**). Notably, several highly dysregulated genes were associated with AD pathology or key oligodendrocyte functions, including *Apbb1* (APP processing), *Lypla2* and *Apoe* (lipid metabolism and transport), *Adamts4* (extracellular matrix remodeling), and *Ndfip1* and *Otulin1* (ubiquitination).

Gene ontology (GO) enrichment analysis highlighted biological processes related to degradation, including “process utilizing autophagic mechanism”, “ubiquitin-dependent protein catabolic process”, and “regulation of cellular catabolic processes” (**Supplementary Fig. 4.2c**). Among the numerous upregulated genes within the “ubiquitin-dependent protein catabolic process” category, *Ubb* and *Fbxw7* were identified, along with other ubiquitin-related E3 ligases (e.g., *Ndfip1*, *Ube3a*), regulatory factors (e.g., *Usp14, Uchl5*), and genes encoding for proteasome subunits (e.g., *Psmd1*, *Psma1*) (**Supplementary Fig. 4.2d**).

These transcriptomic data support the idea that (1) oligodendrocytes are affected early in the course of pathology in AD *in vivo* models, and (2) MYRF accumulation in AD may result from impaired degradation mechanisms, not only *in vitro*, but also *in vivo*.

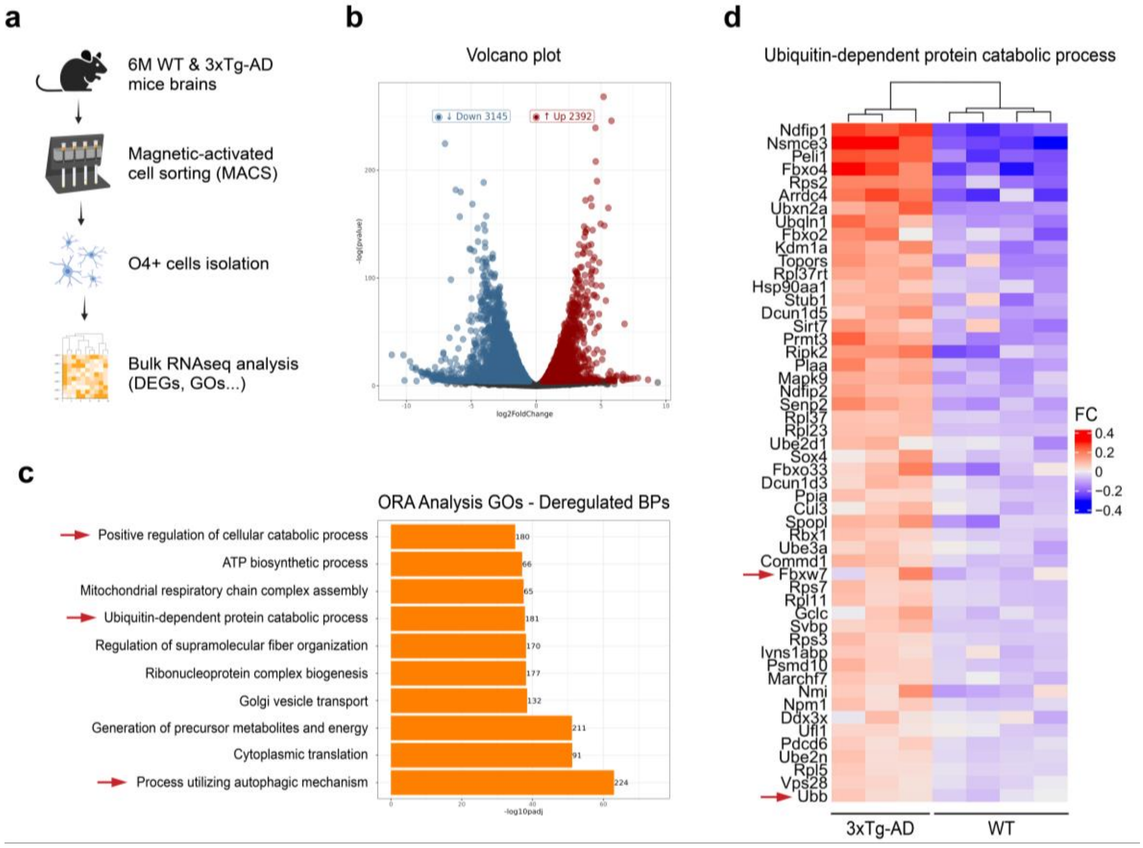


**Supplementary Fig. 4.2.** ***Transcriptomic profiling of MACS-isolated oligodendrocytes reveals alterations in ubiquitin-dependent protein degradation pathways in 3xTg-AD mice.*** (**a**) Schematic of the workflow for MACS-based isolation of $O4^{+}$ oligodendrocytes from brains of 6-month-old WT and 3xTg-AD mice, followed by bulk RNA sequencing (RNA-seq) analysis. $n^{WT} = 4$ and $n^{3xTg\text{-}AD} = 3$ (each n includes 2 pooled brains). (**b**) Volcano plot showing differentially expressed genes (DEGs) in 3xTg-AD versus WT oligodendrocytes. Upregulated DEGs (red) and downregulated (blue). (**c**) Over-representation analysis (ORA) of the top 10 enriched Gene Ontology (GO) biological processes (BP) among DEGs, ranked by gene count. Red arrows highlight GO categories related to protein degradation. (**d**) Heatmap showing relative expression levels of genes within the GO term "Ubiquitin-dependent protein catabolic process" (red = high; blue = low). Genes of interest are indicated by red arrows.

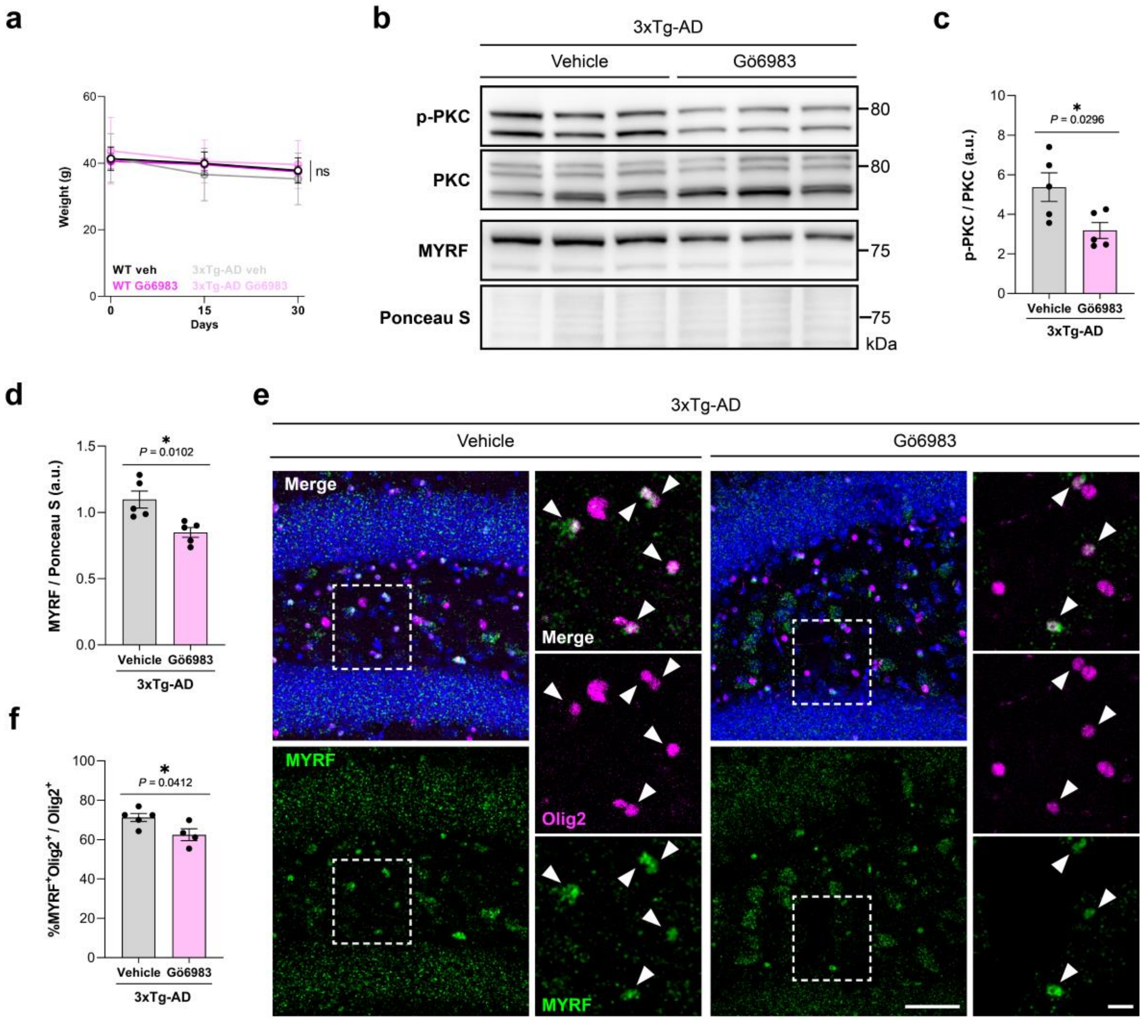


**Supplementary Fig. 5.1.** ***Pharmacological PKC inhibition via intracerebroventricular infusion of Gö6983 reduces MYRF protein levels in the hippocampus of 3xTg-AD mice.*** (**a**) Body weight monitoring of WT and 3xTg-AD mice ICV treated with vehicle or Gö6983 (100nM) throughout the experimental timeline. (**b**) Representative Western blots showing phosphorylated PKC (p-PKC), total PKC, and MYRF levels in contralateral hippocampal lysates from 3xTg-AD mice treated with vehicle or Gö6983. Ponceau S staining was used as a loading control. (**c**) Quantification of PKC activity (p-PKC/PKC ratio) and (**d**) MYRF protein levels normalized to total protein (Ponceau S). (**e**) Representative confocal images of MYRF (green), Olig2 (magenta), and DAPI (blue) immunostaining in the ipsilateral dentate gyrus of ICV treated 3xTg-AD mice. Arrowheads indicate $MYRF^{+}Olig2^{+}$ double-positive OLs. Scale bars: 50 µm (overview), 10 µm (magnification). (**f**) Percentage of $MYRF^{+}$ OLs ($MYRF^{+}Olig2^{+}$) among total $Olig2^{+}$ cells. Data are presented as mean ± S.E.M, with dots representing individual animals. *$p<0.05$; statistical significance was determined by two-tailed unpaired t-test.

**Behavioral assessment of locomotion, anxiety-like behavior, and emotional memory in Gö6983-treated WT and 3xTg-AD mice**

To evaluate spontaneous locomotor and exploratory activity, the OF test was performed on day 1 of the behavioral test battery. In this test, mice were placed in a square arena and allowed to explore freely for 5 minutes. No significant differences were observed across genotypes or treatments in total distance traveled (**Supplementary Fig. 5.2b**), immobility time (**Supplementary Fig. 5.2c**), or anxiety index (**Supplementary Fig. 5.2d**), indicating that gross locomotor activity and basal anxiety-like behavior remain unaffected in 3xTg-AD mice and are not modulated by Gö6983.

On day 2, anxiety-like behavior was further assessed using the EPM, a more sensitive assay for anxiogenic responses. Mice were placed in the center of a plus-shaped maze with two open and two closed arms, and allowed to explore for 5 minutes. No significant differences were found in distance traveled (**Supplementary Fig. 5.2e**) or anxiety index (**Supplementary Fig. 5.2g**). However, a significant increase in immobility time was observed in 3xTg-AD mice compared to WT (**Supplementary Fig. 5.2f**), suggesting a mild hypolocomotion or anxiogenic effect of the genotype; PKC inhibition had no effect.

On day 3, the LD test was used to further assess anxiety-like behavior. Mice were placed in a two-compartment box with one illuminated and one dark chamber, and allowed to explore for 5 minutes. The anxiety index (**Supplementary Fig. 5.2h**) did not differ significantly between groups, indicating no overt anxiogenic phenotype in 3xTg-AD mice and no effect of Gö6983.

To assess emotional learning and memory, we performed the contextual fear conditioning paradigm across three consecutive days, a paradigm largely dependent on amygdala circuits. During acquisition (ACQ), mice received three tone-footshock pairings. All groups showed comparable freezing responses during this phase, indicating similar initial reactivity to the aversive stimulus. However, during the memory retrieval sessions (tone without footshocks) at 24 h (T1) and 48 h (T2), 3xTg-AD mice exhibited significantly increased freezing time compared to WT controls. Gö6983 treatment did not significantly modify freezing behavior in either genotype (**Supplementary Fig. 5.2i**), indicating that PKC inhibition does not alter emotional memory performance.

Collectively, these results confirm that Gö6983 does not affect locomotion or basal anxiety-like behavior, and does not alter the elevated freezing response observed in 3xTg-AD mice, reinforcing the hippocampal specificity of its cognitive effects.

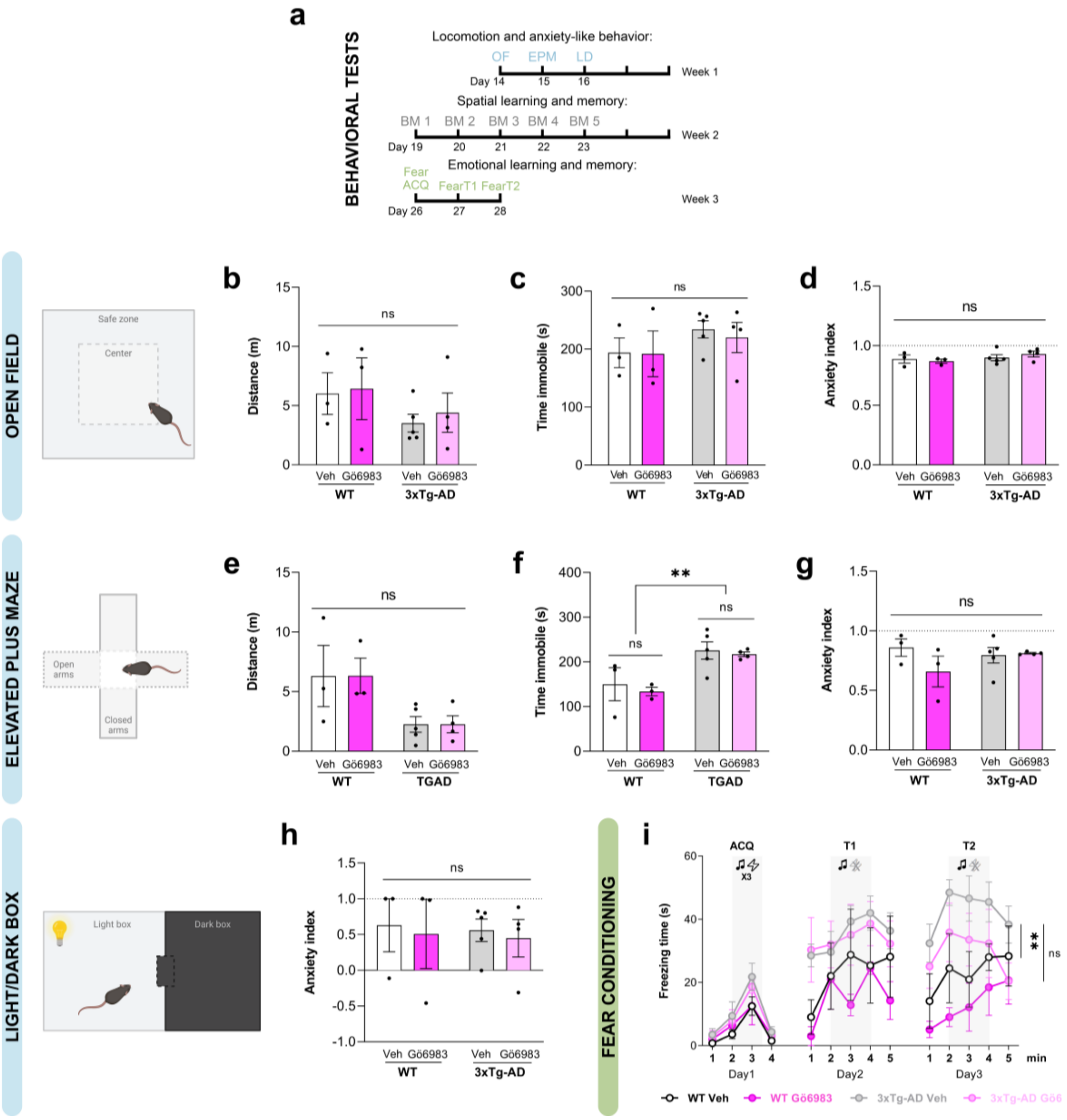


**Supplementary Fig. 5.2.** ***PKC inhibition does not alter innate locomotion, anxiety-like behavior, or emotional memory in 3xTg-AD mice.*** (**a**) Schematic of the behavioral testing schedule: open field (OF), elevated plus maze (EPM), and light/dark box (LD) tests were conducted during week 1 to assess locomotor activity and anxiety-like behavior; Barnes maze (BM) was performed over five consecutive days in week 2 to evaluate spatial learning and memory; and contextual fear conditioning was carried out in week 3 to assess emotional learning and memory. (**b**-**d**) Open field: schematic of the apparatus and quantification of (**b**) distance traveled, (**c**) immobility time, and (**d**) anxiety index (AI). (**e**-**g**) Elevated plus maze: schematic of EPM and quantification of (**e**) distance traveled, (**f**) immobility time, and (**g**) AI. (**h**) Light/dark box: schematic of the LD test and quantification of AI. (**i**) Contextual fear conditioning: quantification of freezing time during acquisition (ACQ; 3 tone–footshock pairings), and after 24 h (T1) and 48 h (T2) (tone but no footshocks). Data are presented as mean ± S.E.M; with dots representing individual animals (n = 3-6 mice per group). **p<0.01, between genotypes; statistical significance was determined by two-way ANOVA followed by Tukey's *post-hoc* test.
